# Come together: Coordinate deconvolution of viral recombination, reassortment, and epistatic survival

**DOI:** 10.64898/2026.11.20.768181

**Authors:** Sergei L. Kosakovsky Pond, Steven Weaver, Danielle Callan, Jordan D. Zehr, Alexander G. Lucaci, Anton Nekrutenko, Darren P. Martin

## Abstract

Viral subtyping and recombination detection underpin taxonomic curation, clinical resistance tracking, and real-time outbreak surveillance. Community nomenclature systems—such as HIV-1 circulating recombinant forms (CRFs), SARS-CoV-2 Pango lineages, and Influenza A genotype catalogs—have created indispensable shared vocabularies, yet rapid archive growth and frequent cross-lineage chimerism now invite a complementary layer that unites speed, provenance, and statistical rigor across viral families. Here, we present `EGGMAN` (Exact Genomic Genealogy and Mosaic Ancestry Nomenclature), a unified framework that separates mapping physical parental coordinates against a principled, non-mosaic founder basis (Tier 1) from statistically guarded assignment to established epidemic clones (Tier 2). By combining reference-free recombination scrubbing (`RhizAeon`), emergence-anchored founder rooting (`ChronAeon`), and portable pathogen specifications, one engine processes contiguous genomes (HIV-1, SARS-CoV-2, wildlife sarbecoviruses) and segmented reassortants (Influenza A) at thousands of sequences per second on a laptop or in a browser—preserving legacy clade names while giving emerging chimeras transparent, coordinate-based identifiers on Day 1. Across public surveillance archives, `EGGMAN` harmonizes the HIV-1 reference catalog and genealogy, reconciles *>* 120,000 clinical *pol* records in Stanford HIVDB, separates emerging CRFs and a monophyletic regional lineage from convergent recombination hotspots, and tracks unannotated H5Nx reassortants and SARS-CoV-2 `X`* waves without temporal lookahead. Finally, census-level mapping across 761 HIV-1 mosaic lineages reveals a two-stage selective sieve separating within-host protein-folding viability (*gp120* and *Gag*-CA cores) from between-host preservation of an intact 3’ regulatory and envelope cassette.

## 1 Introduction

Viral typing and recombination detection serve two foundational roles in virology: organizing genetic diversity into a shared taxonomy and enabling rapid clinical and epidemiological tracking. Across major RNA viruses, community-curated frameworks—including the Los Alamos National Laboratory (LANL) HIV-1 subtype and circulating recombinant form (CRF) compendium [1, 2], SARS-CoV-2 Pango nomenclature [3], Influenza A serotype and genotype systems [4, 5], and clinical drug-resistance databases such as Stanford HIVDB [6]—have made it possible for laboratories worldwide to monitor transmission chains, anticipate drug resistance, and select vaccine strains. Yet RNA viruses do not evolve along strictly bifurcating trees. During cellular co-infection, retroviruses and coronaviruses switch templates to form homologous recombinants [7–11], whereas segmented orthomyxoviruses exchange whole RNA chromosomes to form reassortants [5, 12]. These mosaic events can assemble adaptive mutations across divergent lineages in a single replication cycle, accelerating host-range shifts and immune escape while violating the single-tree assumptions of standard phylogenetic and molecular-clock analyses [13–16].

As public sequence repositories have expanded from hundreds of curated references to tens and hundreds of thousands of full and partial genomes, keeping typing both fast and evolutionarily faithful has become increasingly demanding. Existing computational approaches have addressed complementary halves of this challenge with great success. Dictionary and phylogenetic-placement classifiers such as `COMET` [17], `Nextclade` [18], and `GenoFLU` [4] deliver rapid, standardized tracking of established clades, whereas alignment-based and probabilistic scanners such as `BootScan` [19], `GARD` [14], `SCUEAL` [20], `RDP4/5` [21, 22], and `jpHMM` [23] resolve breakpoints *de novo*. Scaling to contemporary multi-pathogen surveillance, however, highlights five capabilities that would substantially strengthen the field: (i) **principled reference curation** that systematically removes ancestral mosaics from parental alignments [15, 24, 25] while ensuring complete coverage of newly diverged waves; (ii) **transparent, extensible nomenclature** that preserves familiar clinical labels while encoding parental provenance and genome coordinates directly—giving newly emerging chimeras a consistent identifier across laboratories before formal committee naming and linking partial clinical reads [6, 26] cleanly to full genomes; (iii) **cross-pathogen standardization** capable of treating homologous recombination and segment reassortment within a single conceptual grammar; (iv) **explicit statistical rigor** that places each mosaic in phylogenetic and temporal context, filtering out sequencing artifacts and separating true circulating lineages from independent recombination hotspots; and (v) **archive-scale speed**, turning whole-repository recombination mapping from a multi-day cluster task into a routine operation fast enough for census-level evolutionary analysis.

Here, we introduce `EGGMAN` (Exact Genomic Genealogy and Mosaic Ancestry Nomenclature), a unified framework designed around these five goals. Its central architectural principle is to decouple **mapping physical parental ancestry along genome coordinates (Tier 1)** from **testing whether a sequence belongs to an established circulating clone (Tier 2)**. To construct clean, reproducible Tier-1 parental bases without manual curation bias, we pair reference-free recombination detection (`RhizAeon` [27]) with emergence-anchored time-decay rooting (`ChronAeon` [28]) and a formal spanning-basis criterion, storing emergence bounds to support opt-in temporal guarding against retrospective lookahead. Unaligned genomes are projected onto a fixed reference grid and segmented globally within one mathematical objective—using a uniform crossover cost along contiguous genomes and zero cost across RNA segment boundaries—so every sequence receives an invariant, human-readable coordinate formula (`Backbone[Donor:Region]`). In parallel, Tier 2 preserves backward-compatible clade names (`CRF02_AG, XEC, B3.13`) only when a query clears sequential statistical checks for donor coverage, parental dominance, shared post-recombination stem mutations, and absence of secondary third-party tracts. Because genome coordinates, landmarks, and reference panels live in external, hot-swappable pathogen specifications, one native and browser-ready engine deploys across viral families at thousands of sequences per second.

We validate `EGGMAN` across four viral systems—HIV-1 Group M, avian and bovine Influenza A H5Nx, SARS-CoV-2, and wildlife sarbecoviruses—demonstrating that it matches specialized dictionary typers on established clades while delivering profile-HMM–level breakpoint resolution four orders of magnitude faster and recognizing newly emerging chimeras on Day 1. Applying this framework at whole-archive scale yields three broader payoffs: it harmonizes the LANL HIV-1 reference compendium and organizes its CRFs into a three-generation genealogy; it reconciles 120,468 clinical *pol* records in Stanford HIVDB and separates emerging CRFs and a monophyletic regional lineage from convergent hotspots across *>* 28,500 GenBank hosts; and it turns census-level HIV-1 mosaic maps into a quantitative evolutionary assay, revealing a two-stage selective sieve that separates within-host protein-folding viability from between-host preservation of an intact 3’ regulatory and envelope cassette.

## 2 Materials and Methods

### 2.1 Conceptual overview: separating ancestry mapping from clade naming

Viral classification combines two distinct evolutionary questions that are easiest to solve when kept separate: (1) *What parental lineages make up this genome, and where are the junctions between them?* and (2) *Does this isolate belong to an established, clonally spreading epidemic lineage?* Combining both tasks into a single flat reference dictionary creates a structural tension: omitting a major clonal clade turns its members into false mosaics of neighboring references, whereas retaining an unmasked recombinant inside the parental basis propagates hidden bias into every downstream breakpoint call and molecular-clock estimate.

`EGGMAN` resolves this tension with a two-tier architecture parameterized entirely through external, human-readable `PathogenSpec` packs (`--pack`; Table 1). Every query sequence is projected onto a canonical pathogen coordinate system and evaluated in a single fast pass across two complementary layers (Fig. 1A):

- **Tier 1 (Ancestry Basis)** holds a compact set of strictly non-mosaic basal founders used to decompose any genome—full or partial, known or completely novel—into an invariant genomic coordinate map.
- **Tier 2 (Epidemic Catalog)** holds curated profiles of circulating epidemic lineages (such as HIV-1 CRFs, Influenza A reassortant genotypes, or SARS-CoV-2 X* recombinants) and tests whether a query genuinely descends from one of those named clones.

**Figure 1:**
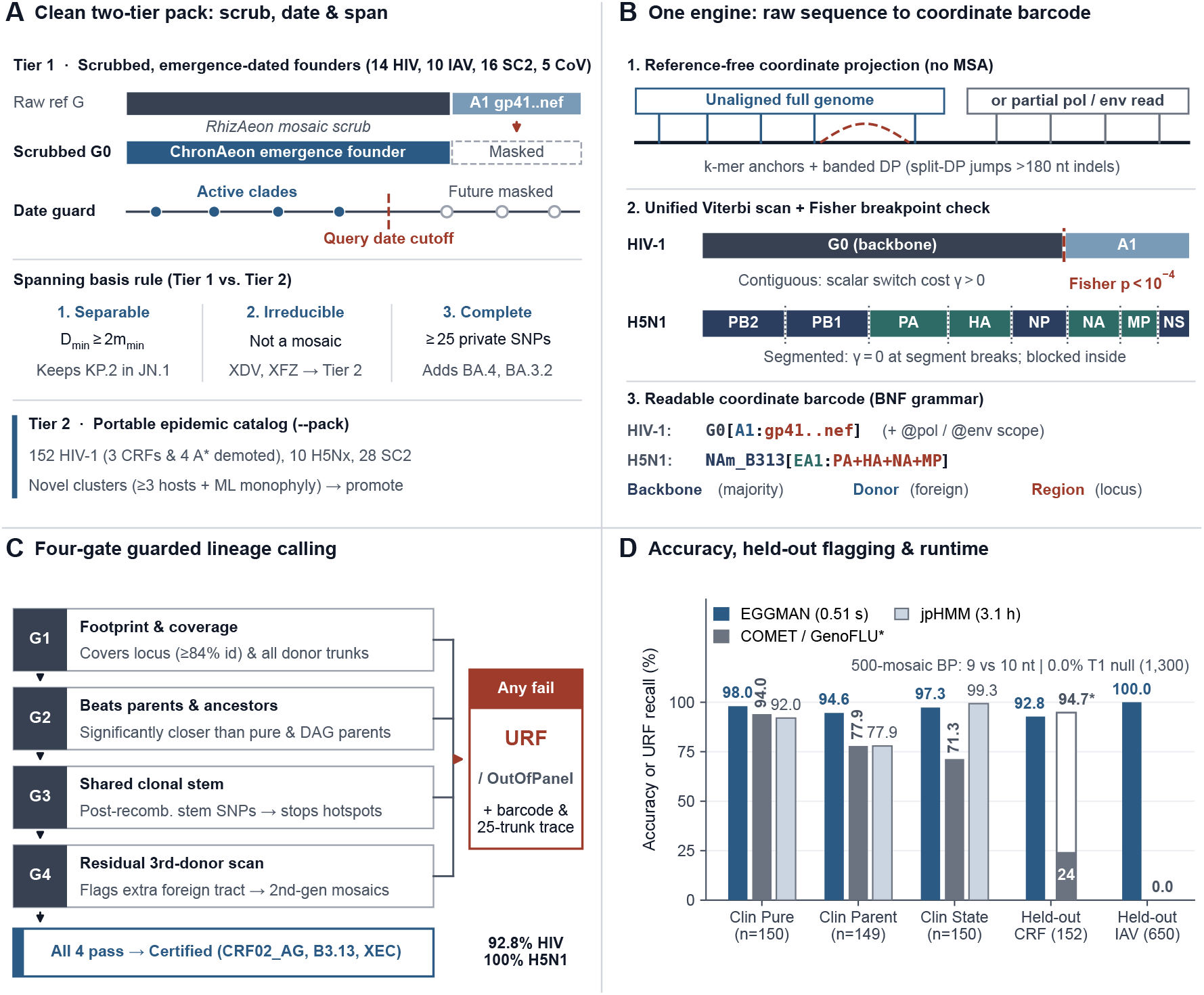
EGGMAN two-tier pack construction, unified segmentation, four-gate certification, and out-of-sample HIV-1 benchmark performance. **A:** Tier 1 stores strictly non-mosaic founders scrubbed via `RhizAeon` (e.g., masking `A1:gp41..nef` inside Subtype G to yield `G0`) and emergence-weighted via `ChronAeon` with an opt-in temporal guard masking post-sample clades, while the three-criterion Spanning Basis Rule separates divergent clonal waves (Tier 1) from circulating mosaics in the portable Tier-2 catalog. **B:** Unaligned full genomes or partial amplicons project onto canonical coordinates without multiple sequence alignment, pass through one discrete Viterbi objective (constant *γ >* 0 for contiguous recombination; *γ* = 0 at segment boundaries for Influenza A reassortment) checked by exact Fisher tests, and render into readable BNF barcodes (`Backbone[Donor:Region]`). **C:** Four sequential certification gates test foot-print coverage, parental/DAG dominance, shared post-recombination stem mutations (stopping convergent hotspots such as D[A1:env]), and residual third-donor tracts (catching second-generation mosaics), safely routing failures to `URF` (92.8% `URF` + 2.0% OutOfPanel = 94.7% non-legacy on held-out HIV-1; 100% H5N1). **D:** Out-of-sample accuracy, independent LANL parent-set recovery, held-out novelty flagging, and runtime across clinical HIV-1 cohorts, 500-mosaic synthetic breakpoints, and 1,300 null simulations versus `COMET` v2.4, `jpHMM`, and `GenoFLU` (*`COMET` 94.7% total unassigned comprises 24% hint-free [solid] plus 70% carrying legacy CRF hints [open]; `GenoFLU` returns 0% novel coordinate recovery on held-out H5N1).

**Table 1:**
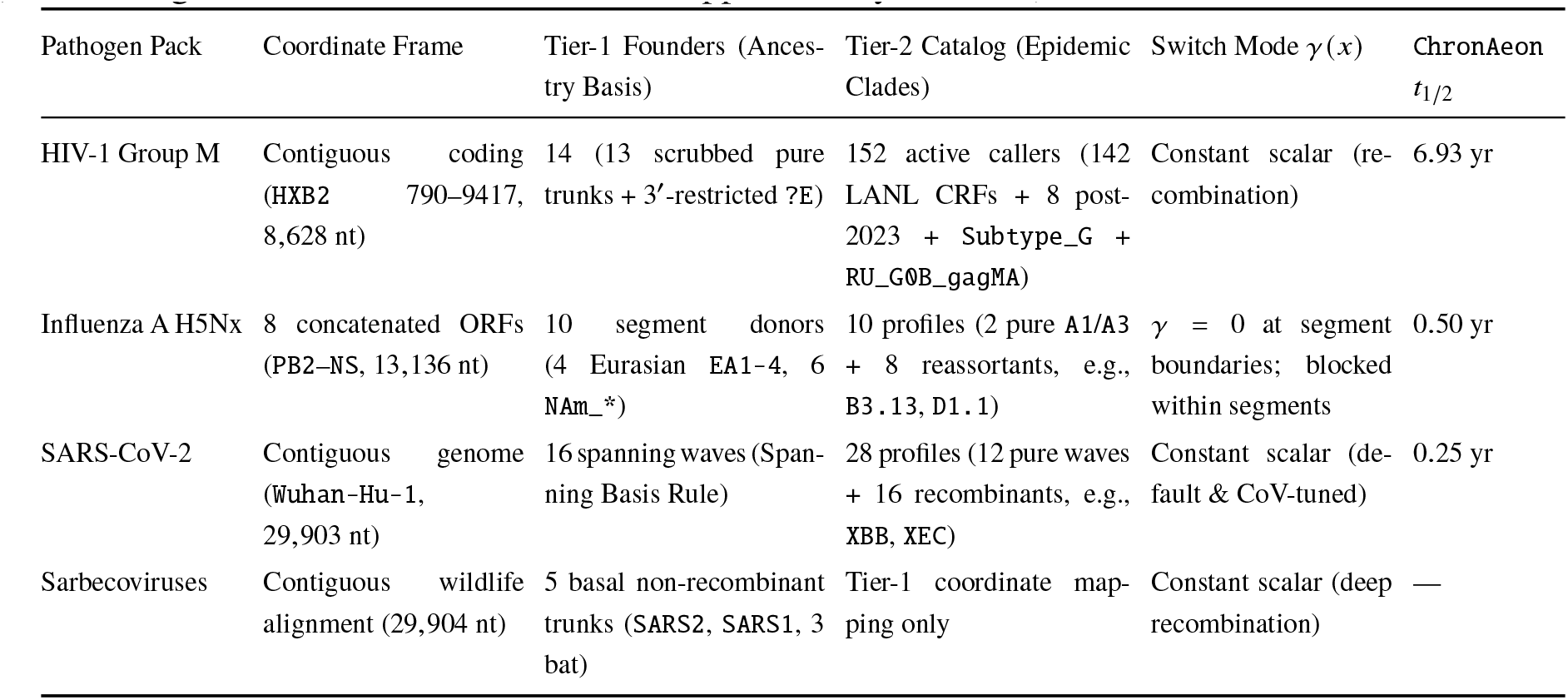
Summary of external PathogenSpec reference packs evaluated with the unified EGGMAN engine (detailed alignment and Viterbi constants in Supplementary Table S1).

| Pathogen Pack | Coordinate Frame | Tier-1 Founders (Ances-try Basis) | Tier-2 Catalog (Epidemic Clades) | Switch Mode γ(x) | ChronAeon t<sub>1/2</sub> |
|---|---|---|---|---|---|
| HIV-1 Group M | Contiguous coding (HXB2 790–9417, 8,628 nt) | 14 (13 scrubbed pure trunks + 3′-restricted ?E) | 152 active callers (142 LANL CRFs + 8 post-2023 + Subtype_G + RU_G0B_gagMA) | Constant scalar (re-combination) | 6.93 yr |
| Influenza A H5Nx | 8 concatenated ORFs (PB2–NS, 13,136 nt) | 10 segment donors (4 Eurasian EA1-4, 6 NAm_*) | 10 profiles (2 pure A1/A3 + 8 reassortants, e.g., B3.13, D1.1) | γ = 0 at segment boundaries; blocked within segments | 0.50 yr |
| SARS-CoV-2 | Contiguous genome (Wuhan-Hu-1, 29,903 nt) | 16 spanning waves (Span-ning Basis Rule) | 28 profiles (12 pure waves + 16 recombinants, e.g., XBB, XEC) | Constant scalar (de-fault & CoV-tuned) | 0.25 yr |
| Sarbecoviruses | Contiguous wildlife alignment (29,904 nt) | 5 basal non-recombinant trunks (SARS2, SARS1, 3 bat) | Tier-1 coordinate map-ping only | Constant scalar (deep recombination) | — |

Because all viral coordinates, parental profiles, and lineage catalogs live outside the compiled engine, the identical algorithm operates across retroviruses, segmented orthomyxoviruses, and coronaviruses without changing a line of core code.

### 2.2 Building a clean, self-consistent reference pack

Constructing a trustworthy `PathogenSpec` pack relies on three principled curation steps that ensure reference purity, chronological fidelity, and completeness as epidemics evolve (Fig. 1A; exact equations, coordinates, and weights in Supplementary Methods S1.1–S1.2 and Table S1):

1. **Purging ancestral mosaics (**`RhizAeon`). Before freezing Tier-1 founders, candidate pure clades are screened using reference-free recombination detection (`RhizAeon` [27]) to locate and mask ancestral chimeric tracts rather than letting them masquerade as pure background. In HIV-1, this step masks the ancestral Subtype A1 envelope patch (*gp41–nef*) embedded inside canonical Subtype G references—yielding the clean basal trunk `G0`—masks a 3’`K:nef` segment in Subtype L, and restricts the extinct Asian `CRF01_AE` donor `?E` to the 3’ half of the genome (*vif–nef*) where it actually circulates. The resulting HIV-1 Tier-1 basis consists of 14 clean founders (13 Group M trunks plus 3’-restricted `?E`), while non-HIV packs comprise 10 Influenza A H5Nx segment donors (four Eurasian and six North American), 16 SARS-CoV-2 spanning waves, and 5 non-recombinant bat/human sarbecovirus trunks (Table 1).
2. **Emergence-anchored profiles and non-lookahead timing (**`ChronAeon`). Standard majority-rule consensus sequences drift toward whichever recent outbreak happens to be most heavily sequenced. Instead, `EGGMAN` weights each founder alignment exponentially toward its earliest emergence using deletion-aware `ChronAeon` time-decay profiling [28] (*w_i_* ∝ 2^−(^*^ti^* ^−^*^t^*^0)/^*^t^*^1/2^, with half-lives *t*_1/2_ = 6.93 yr for HIV-1, 0.50 yr for Influenza A, and 0.25 yr for SARS-CoV-2). Without tree search, `ChronAeon` also estimates each clade’s emergence time (*t*_MRCA_) and uncertainty bounds, validated against IQ-TREE 2 [29] phylogenies dated with `LSD2` [30] and `TreeTime` [31]. Storing each clade’s emergence floor *t*_floor_(*k*) enables an opt-in temporal guard (`--temporal-guard` with `--temporal-mode sampled| submission`) that admits donor or catalog clade *k* only when *t*_floor_(*k*) ≤ *t*_query_ + *δ_t_* —using earliest collection dates (*t*_MRCA_ fallback) with pack-calibrated slack *δ_t_*(3.0 yr HIV-1, 0.5 yr non-HIV) in sampled mode, or earliest public deposition dates with *δ_t_*= 0 in submission mode (falling back to earliest sampled dates on HIV-1 basal trunks) to prevent retrospective lookahead bias during surveillance audits.
3. **Systematic completeness without bloat (Spanning Basis Rule).** During fast-moving outbreaks such as SARS-CoV-2, new divergent clonal waves periodically arise. To decide objectively when a candidate wave *c* belongs in the Tier-1 basis B versus Tier 2, we apply a three-criterion Spanning Basis Selection Rule evaluated strictly on early representatives (≤ 25 genomes per candidate): (a) *separability*, requiring pairwise coding distance *D*_min_(*c,* B) ≥ 2*m*_min_ from every existing Tier-1 founder to avoid splitting shallow sublineages (e.g., keeping `KP.2` inside `JN.1`); (b) *irreducibility*, requiring that screening *c* against B produces no statistically supported mosaic decomposition (automatically routing recombinants such as `XDV` and `XFZ` to Tier 2); and (c) *completeness*, promoting any non-mosaic wave that would otherwise induce *>* 50% pseudo-mosaic calls among its members or carries saltation-scale private divergence (≥ 25 coding substitutions). Applying this rule to an initial 11-wave SARS-CoV-2 basis cleanly promoted five divergent waves (`P.1, BA.4, BA.3.2, LF.7`, and `LP.8.1`) to establish the frozen 16-founder spanning basis without overfitting to test cohorts.

### 2.3 From raw sequence to coordinate barcode (Tier 1)

Turning an unaligned viral genome—or a short clinical amplicon—into an invariant ancestry map happens in three fast, deterministic stages without building a multiple sequence alignment (Fig. 1B; full alignment, split-DP, and Viterbi equations in Supplementary Methods S1.3 and Table S1):

1. **Reference-free coordinate projection.** Each input FASTA sequence is mapped directly onto the pathogen’s canonical coordinate grid by matching exact 15-mer seeds (step 2) against all indexed reference profiles on both forward and reverse-complement strands, selecting the higher-scoring orientation (with an 11-mer fallback when forward/reverse 15-mer hits are sparse and local 11-mer rescue across *>* 350 nt unanchored gaps). Collinear hits are chained into anchors and intervening spans are aligned by banded Needleman–Wunsch (+2 match, –1 mismatch, –3 linear gap; band 45) against a 4-bit A/C/G/T reference-profile union mask (query ambiguity codes are ignored during subsequent ancestry scoring). Whenever flanking anchors reveal a large structural insertion or deletion (|*m* – *n*| *>* 180 nt), a two-piece split dynamic-programming bridge (band 45) jumps cleanly across the indel rather than smearing bases across neighboring genes. Full-length genomes and partial clinical reads (*pol*, *env*, or Spike) thus land on the exact same numbered coordinate frame in milliseconds.
2. **One unified segmentation model for recombination and reassortment.** Across the aligned coding span *x* ∈ {1*,…, L*}, a discrete Viterbi dynamic program assigns each site a state *s_x_* ∈ S drawn from the Tier-1 founders plus an explicit `Unattributed` ghost state that absorbs divergent, out-of-panel sequence rather than forcing it into false mosaics:

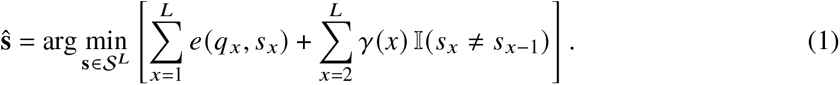 The emission cost *e*(*q_x_, s_x_*) is defined in mismatch-equivalent (Hamming) units: 0 for an exact A/C/G/T match, 1.0 for a mismatch, 0.35 for a gap or ambiguous reference site, 0.58 for the `Unattributed` ghost state (acting as an out-of-panel divergence and alignment-failure floor above 58% mismatch), and 2.0 outside a domain-restricted trunk (such as `?E` upstream of `HXB2` 5097, plus a 0.035/site prior inside its 3’ domain); query gaps and IUPAC ambiguity symbols contribute 0. Under a symmetric substitution probability *p*, one mismatch corresponds to a log-likelihood ratio of ln[(1 – *p*)/(*p*/3)] nats, so a mismatch-unit switch penalty *γ* corresponds to a steeper log-likelihood barrier at low divergence (*p* ≈ 10^−3^ in SARS-CoV-2) than at high divergence (*p* ≈ 0.08 in HIV-1)—directly explaining why low-divergence coronaviruses use a smaller mismatch-unit switch penalty (*γ* = 3.5) than HIV-1 (*γ* = 7.0). Crucially, the position-dependent switch penalty *γ* (*x*) unifies homologous recombination and viral reassortment inside Eq. (1): on contiguous genomes (HIV-1, SARS-CoV-2, sarbecoviruses), *γ* (*x*) ≡ *γ* is a constant divergence-calibrated penalty (default *γ* = 7.0, with tunable presets for compendium auditing, screening, and low-divergence coronavirus tracking; Supplementary Table S1); on segmented genomes (Influenza A H5Nx), *γ* (*x*) = 0 at inter-segment boundaries—allowing free reassortment across the eight RNA segments—while *γ* (*x*) = 10^6^ inside segments whenever intra-segment recombination is disallowed.
3. **Statistical breakpoint falsification and coordinate barcode grammar.** Dynamic programming paths alone can occasionally chase localized sequencing noise or convergent point mutations. `EGGMAN` therefore tests every proposed junction between adjacent states *A* and *B* against three lightweight guards before accepting it: each flanking segment must span at least *L*_min_ reference bases (120 nt full, 150 nt partial, or half-segment on segmented packs), carry at least *m*_min_ parent-discriminating substitutions (5 default; 3 CoV-tuned), and exhibit significant parental switching across the boundary under an upper-tail 2 × 2 Fisher’s exact test on left/right diagnostic counts (*n_L, A_, n_L,B_*; *n_R, A_, n_R,B_*) (*p <* 10^−4^ default). Unsupported switches collapse back into the surrounding backbone. Verified segment maps render directly into a deterministic coordinate barcode governed by a formal Backus–Naur Form grammar (`<backbone>(Clonal) or <backbone>[<donor>:<span_list>`]; Supplementary Table S2). The Tier-1 founder spanning the largest fraction of aligned sequence serves as the backbone, and each verified foreign insertion or reassorted segment appears in brackets by donor and canonical landmark—yielding self-documenting Day-1 identifiers across viral families, such as G0[A1:gp41..nef] in `HIV-1, NAm_B313[EA1:PA+HA+NA+MP]` in bovine H5N1 (B3.13), or `Bat_HKU3[SARS1_HSZ:S+N..ORF10]` in wildlife sarbecoviruses. Partial clinical amplicons carry an explicit locus suffix (e.g., @pol or @env), linking partial diagnostic reads transparently to matching full-genome mosaics.

### 2.4 Certifying epidemic lineages versus novel recombinants (Tier 2)

While Tier 1 gives every sequence a transparent coordinate barcode, clinical and public-health surveillance also depends on preserving trusted lineage names (`CRF02_AG, B3.13, XEC`). Tier 2 bridges both worlds by testing each aligned query against the pack’s audited catalog of circulating epidemic clades (Table 1; complete gate equations and SNP/*p*-value cutoffs in Supplementary Methods S1.4 and Table S1). Crucially, assigning the closest catalog profile by raw similarity alone risks two systematic errors: mislabeling an independent convergent recombinant that breaks at the same genomic hotspot as a known CRF, or hiding a new second-generation mosaic under its parent CRF’s name.

To guard against both failure modes, every candidate epidemic lineage *c* must clear four sequential statistical falsification gates (Fig. 1C):

1. **Footprint and donor coverage (Gate 1).** *Does the query span the expected locus at suflcient identity and show evidence from every constituent parental donor of c?* Gate 1 requires ≥ 100 nt aligned overlap, ≥ 84% sequence identity to *c*, coverage of at least two donor blocks (≥ 120 nt each), and donor plurality (*n*_donor_ *> n*_backbone_ on ≥ 3 diagnostic SNPs) across every constituent donor block.
2. **Parental and ancestral dominance (Gate 2).** Does the query match c significantly better than every pure parental trunk and every immediate ancestor p in the recombination genealogy (DAG)? Across all sites where *c* and *p* differ, Gate 2 requires *n_c_ > n_p_* with excess Δ_2_ = *n_c_* – *n_p_* ≥ *d*_min_ and one-sided binomial support *P*(Bin(*n_c_* + *n_p_,* 0.5) ≥ *n_c_*) *< α*_2_ (defaults *d*_min_ = 5 full / 3 partial, *α*_2_ = 10^−3^ / 10^−2^).
3. **Shared clonal history (Gate 3—stem test).** *Does the query carry the private “stem” mutations that accumulated after the founding recombination event?* Constructing the composite parental expectation *B_c_* (*x*) by stitching each pure parent’s profile along *c*’s canonical barcode, Gate 3 counts query matches to *c* versus *B_c_* at every site where *c*(*x*) ≠ *B_c_* (*x*) and requires net stem excess Δ_stem_ = *s_c_* – *s_B_* ≥ *s*_min_ with one-sided binomial *p < α*_3_ (pack-specific defaults in Supplementary Table S1)—separating clonal descendants of a single founder event from independent convergent chimeras at the same hotspot.
4. **Residual scan for hidden secondary tracts (Gate 4).** After accounting for c, is there any localized stretch of leftover mismatches better explained by a third competing clade k? Scoring each informative site by *r_x_* = +1 where the query matches *k* over *c* and –1 where it matches *c* over *k*, an all-excursion Kadane scan 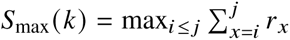 flags any ≥ 80 nt window exceeding the net advantage floor (≥ 12 full / ≥ 10 partial, Fisher *p <* 10^−4^), catching second-generation mosaics before they can be misfiled under their parent label.

Sequences failing any gate are never forced into a legacy name. Instead, they fall back safely to `URF` while keeping their invariant Tier-1 coordinate barcode. Separately, if any Tier-1 segment (≥ 80 valid nt) falls below the pack’s minimum segment identity floor (82% HIV-1, 90% Influenza A and SARS-CoV-2, 70% sarbecoviruses) or violates an active temporal guard floor, that segment is relabeled OutOfPanel and the sequence routes immediately to `OutOfPanel` (while fully `Unattributed` single-segment queries return `Unclassifiable`). For HIV-1, `URF` outputs are paired with an interpretable 25-trunk epidemiological trace (combining the 13 pure basal trunks excluding `?E` with 12 major pandemic founder CRFs, plus `Unattributed`) so second- and third-generation mosaics immediately display their epidemic backbone (such as `CRF01_AE, CRF02_AG`, or `CRF07_BC`). Finally, in our post-2021 SARS-CoV-2 streaming surveillance scripts, a lightweight post-processing quality-assurance filter flags localized Wuhan-B amplicon-backfill artifacts—where primer dropout causes consensus pipelines to splice reference sequence into an Omicron genome and mimic recombination.

### 2.5 Cohorts, benchmarking, stress tests, and orthogonal validation

To evaluate accuracy, speed, and real-world surveillance utility without circularity, we organized our validation around four complementary pillars (detailed dataset inventories, comparator commands, and benchmark breakdowns in Table 2, Supplementary Methods S1.5, and Supplementary Tables S3–S10):

1. **Public surveillance archives across four viral systems.** *HIV-1:* We audited the 2023 LANL Reference and filtered full-genome alignments (*N* = 518 and 5,088), NCBI GenBank full-length genomes (31,825 sequences deduplicated at *<* 0.8% divergence within identical barcodes into 7,720 independent patients) plus strictly partial-only GenBank and LANL *pol* and *env* records (38,250 sequences across 20,797 patients), and the Stanford HIVDB [6] reference panel (*N* = 720) and clinical surveillance archive (*N* = 120,468, separated into 111,905 `isolates`.d-linked and 8,563 unlinked records). *Influenza A, SARS-CoV-2, and wildlife sarbecoviruses:* Cross-family evaluation spanned 1,200 complete eight-segment SRA H5N1 genomes and 3,980 complete BV-BRC H5Nx assemblies (2020–2026); 1,000 curated clinical SARS-CoV-2 genomes, 10,062 Global World Union genomes, and two live BV-BRC streams (23,376 stratified and 100,000 genomes containing 77,358 from 2025–2026); and 79 valid whole-genome bat and pangolin sarbecoviruses (excluding four all-N placeholders).
2. **Head-to-head comparisons against specialist tools.** We benchmarked `EGGMAN` directly against the standard reference classifier for each viral system: `COMET` v2.4 [17] and 28-worker `jpHMM` [23] across 1,093 unaligned HIV-1 queries spanning 12 curated reference, clinical *pol*/*env*, CRF, and `URF` panels; 4-thread USDA `GenoFLU` [4] across complete eight-segment H5N1 genomes; and 8-thread `Nextclade` CLI [18] across curated SARS-CoV-2 waves and recombinants.
3. **Controlled stress tests for breakpoints, specificity, novelty, and non-lookahead timing.** *Known-breakpoint accuracy:* Exact nucleotide localization was measured across 500 simulated GTR + Г_4_ mosaic HIV-1 genomes containing 900 true crossover junctions. *False-positive calibration:* We simulated 1,300 strictly non-recombinant HIV-1 genomes across all 13 pure founders under standard evolution, 3× rate acceleration, and APOBEC3G/3F hypermutation. *Held-out novelty flagging:* We tested safe `URF` fallback via leave-one-sequence-out and lineage-omitted HIV-1 cross-validation (*N* = 536), matched CRF01-118 masking (*N* = 152), withheld bovine/avian H5N1 B3.13/D1.1 holdouts (*N* = 650), and held-out SARS-CoV-2 `XDV`/`XFZ`/XFC genomes. *Clock and temporal-guard validation:* `ChronAeon` emergence dates (95% block-bootstrap and Fieller confidence intervals [32]) were benchmarked across 21 HIV-1 clades (3,594 genomes), 32 SARS-CoV-2 clades, and 15 Influenza A alignments against `IQ-TREE` 2 phylogenies dated with `LSD2` and `TreeTime`, and `--temporal-guard` masking was verified across 16 collection-date and submission-date ablations.
4. **Orthogonal phylogenetic verification.** Every surveillance finding surfaced by coordinate mapping—including candidate regional HIV-1 lineages and convergent hotspots in GenBank, 40 West African `CRF02_AG` clinical *pol* controls, 15 USDA-unassigned H5N1 reassortants (16 segment trees), BV-BRC H5Nx genotypes, and unannotated SARS-CoV-2 `URF` clusters—was tested independently by segment-level maximum-likelihood tree reconstruction in `IQ-TREE 2` (GTR + Г_4_, 1,000 `UFBoot2` replicates [29, 33]).

**Table 2:**
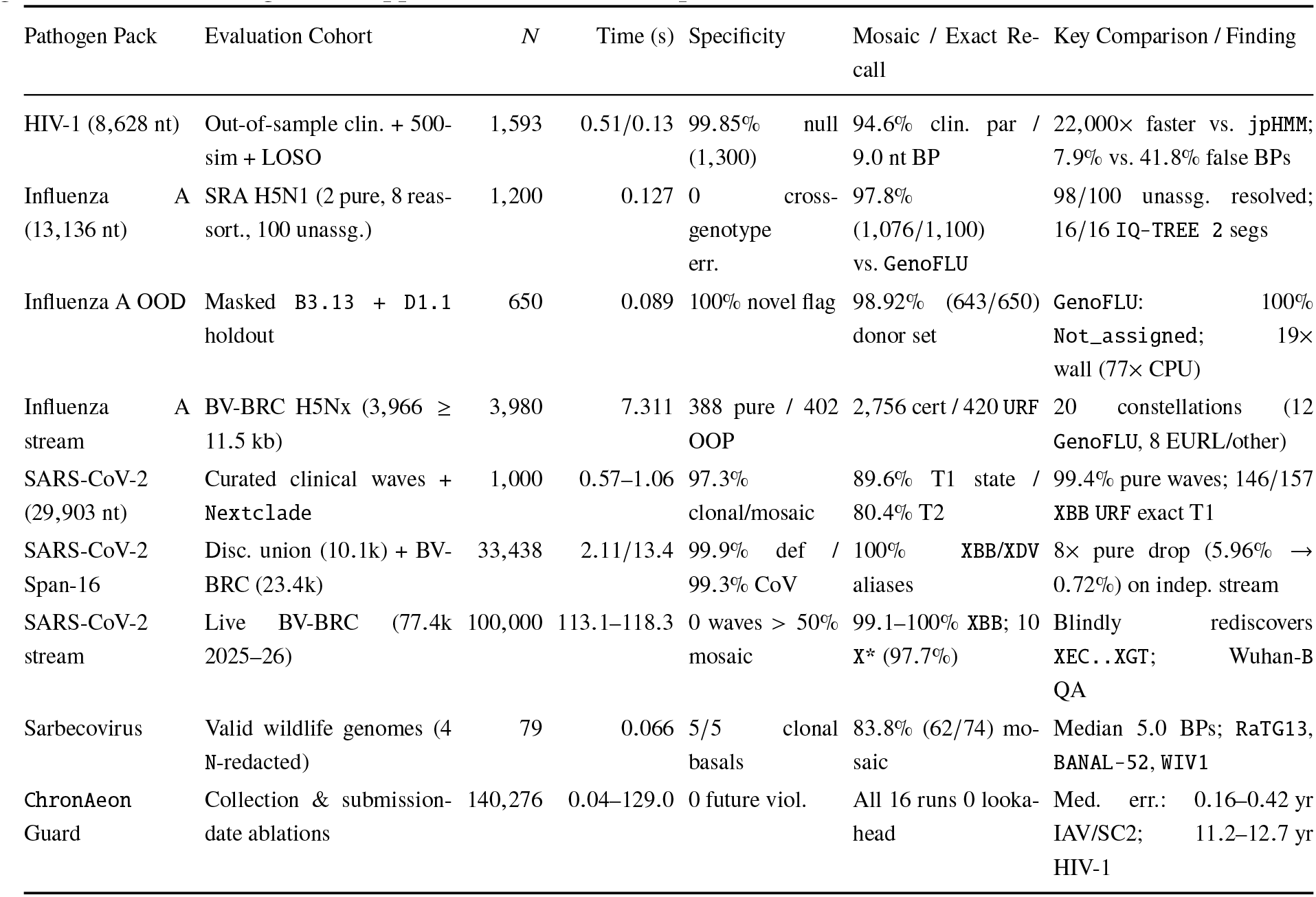
Multi-pathogen EGGMAN performance, external comparator benchmarks, and ChronAeon temporal-guard validation using hot-swappable PathogenSpec panels (--pack).

| Pathogen Pack | Evaluation Cohort | *N* | Time (s) | Specificity | Mosaic / Exact Re-call | Key Comparison / Finding |
|---|---|---|---|---|---|---|
| HIV-1 (8,628 nt) | Out-of-sample clin. + 500-sim + LOSO | 1,593 | 0.51/0.13 | 99.85% null (1,300) | 94.6% clin. par / 9.0 nt BP | 22,000× faster vs. jpHMM; 7.9% vs. 41.8% false BPs |
| Influenza A (13,136 nt) | SRA H5N1 (2 pure, 8 reas-sort., 100 unassg.) | 1,200 | 0.127 | 0 cross-genotype err. | 97.8% (1,076/1,100) vs. GenoFLU | 98/100 unassg. resolved; 16/16 IQ-TREE 2 segs |
| Influenza A OOD | Masked B3.13 + D1.1 holdout | 650 | 0.089 | 100% novel flag | 98.92% (643/650) donor set | GenoFLU: 100% Not_assigned; 19× wall (77× CPU) |
| Influenza A stream | BV-BRC H5Nx (3,966 ≥ 11.5 kb) | 3,980 | 7.311 | 388 pure / 402 OOP | 2,756 cert / 420 URF | 20 constellations (12 GenoFLU, 8 EURL/other) |
| SARS-CoV-2 (29,903 nt) | Curated clinical waves + Nextclade | 1,000 | 0.57–1.06 | 97.3% clonal/mosaic | 89.6% T1 state / 80.4% T2 | 99.4% pure waves; 146/157 XBB URF exact T1 |
| SARS-CoV-2 Span-16 | Disc. union (10.1k) + BV-BRC (23.4k) | 33,438 | 2.11/13.4 | 99.9% def / 99.3% CoV | 100% XBB/XDV aliases | 8× pure drop (5.96% → 0.72%) on indep. stream |
| SARS-CoV-2 stream | Live BV-BRC (77.4k 2025–26) | 100,000 | 113.1–118.3 | 0 waves > 50% mosaic | 99.1–100% XBB; 10 X* (97.7%) | Blindly rediscovers XEC..XGT; Wuhan-B QA |
| Sarbecovirus | Valid wildlife genomes (4 N-redacted) | 79 | 0.066 | 5/5 clonal basals | 83.8% (62/74) mo-saic | Median 5.0 BPs; RaTG13, BANAL-52, WIV1 |
| ChronAeon Guard | Collection & submission-date ablations | 140,276 | 0.04–129.0 | 0 future viol. | All 16 runs 0 looka-head | Med. err.: 0.16–0.42 yr IAV/SC2; 11.2–12.7 yr HIV-1 |

### 2.6 Census-level HIV-1 supply and two-stage selective sieve analysis

High-throughput coordinate mapping transforms public viral archives into a quantitative assay of recombinant emergence and natural selection. Combining deduplicated full-length GenBank HIV-1 genomes with active CRFs, we tested ecological supply, epidemic persistence, and structural selection across three linked statistical steps (exact locus coordinates and test equations in Supplementary Methods S1.6 and Table S9):

1. **National co-infection supply versus *de novo* `URF` emergence.** Across *R* = 32 countries represented by at least 40 deduplicated hosts, we quantified local circulating lineage diversity using non-URF heterozygosity 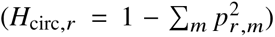 and tested whether higher subtype co-circulation predicts higher *de novo* `URF` prevalence. Associations were assessed with Spearman rank correlation, sample-size–weighted least squares (WLS), heteroskedasticity-consistent binomial regression (`HC3`), and offset negative-binomial models controlling for Sub-Saharan African geography, host sample size, and sampling span.
2. **Heavy-tailed cluster scaling and deduplication sensitivity.** To measure how steeply recombinant lineages drop off between single-host emergence and sustained transmission, we fit observed patient cluster sizes across all *N* = 761 distinct HIV-1 mosaic lineages by maximum likelihood to discrete Hurwitz-zeta (*P*(*N* = *k*) = *k*^−^*^α^*/*ζ* (*α, x*_min_)), discrete lognormal, Yule–Simon, and negative-binomial distributions, evaluating power-law goodness of fit via Clauset bootstrap testing and Vuong likelihood-ratio comparisons [34], and testing invariance of cluster proportions and Stage-2 rates across host deduplication thresholds (0.4%, 0.8%, and 1.5% Hamming divergence).
3. **Two-stage selective sieve (within-host folding viability versus between-host transmission).** To separate constraints required merely to assemble a viable virion inside one patient from those favoring epidemic spread across hosts, we compared mosaic architectures across three nested survival tiers: 491 strictly singleton dead-end `URF`s (2,046 breakpoints), 151 established CRFs (701 breakpoints), and 22 major CRFs circulating in ≥ 10 hosts (93 breakpoints). *Stage 1—Within-host protein-folding filter:* We tested breakpoint depletion inside rigid protein-folding cores (*gp120* core and *Gag*-CA capsid core, alongside *Pol*-RT and canonical full-gene boundaries) and enrichment inside four flexible inter-domain hinges against three nested null expectations: (a) uniform coding span, (b) empirical parent-pair–weighted diagnostic-SNP density (masking hypervariable *env* loops), and (c) explicit `EGGMAN` detectability nulls constructed by recalling both 1,000 simulated single-crossover mosaics (2% noise; 1,204–1,222 recalled breakpoints) and 1,000 realistic 4-crossover mosaics at 6%–10% GTR + Г_4_ divergence (4,000 true breakpoints; 4,126 recalled) through the identical inference pipeline. Matching observed depletion against both detectability nulls rules out methodological blind spots in breakpoint calling. *Stage 2—Between-host transmission filter:* Comparing established and major CRFs directly against singleton `URF`s, we evaluated topological remodeling and multi-locus co-ancestry using 10,000-replicate linear segment permutations, circular genome shifts, parental-trunk–stratified Cochran–Mantel–Haenszel (CMH) odds ratios, multivariable logistic regression (adjusting for total breakpoint count and parental trunk pair), and Fisher exact tests. Crucially, to eliminate phylogenetic pseudoreplication from Generation 2–3 CRF hubs, we repeated every Stage-2 test after restricting established CRFs to the 71 alias-collapsed Generation-1 primary founder CRFs and after deduplicating inherited DAG junctions (527 *de novo* CRF breakpoints)—contrasting isolated point exon linkage (*tat1* ↔ *tat2*) against preservation of the full undivided 3’ regulatory and envelope cassette (`HXB2` 5831–8653: *tat1–vpu–env–tat2–rev2*), *RRE* ↔ *rev2* linkage, upstream *gag/pol* shifts, and *gp41*-CT return crossovers.

## 3 Results

### 3.1 Two-tier separation preserves speed and precision across hard HIV-1 regimes where flat classifiers diverge

We evaluated `EGGMAN` across 1,093 unaligned HIV-1 genomes using independent published LANL and GenBank subtype/CRF annotations as the reference standard for empirical cohorts, and known crossover coordinates and pure founders for synthetic tests (Fig. 1D; complete per-panel metrics in Table 2 and Supplementary Tables S4–S6). Separating out-of-sample generalization from in-sample catalog consistency clarifies both where tools agree and where two-tier architecture separates them. On out-of-sample clinical pure subtypes (*n* = 150), all comparators performed strongly, with `EGGMAN` achieving 98.0% exact accuracy versus 94.0% for `COMET` v2.4 and 92.0% for `jpHMM` while running at order 10^3^ seq/s in native Rust—processing the entire 1,093-genome benchmark in under a second locally, four orders of magnitude faster than 28-worker `jpHMM` (*>* 3 h wall time). In-sample catalog consistency on the 142 active LANL CRF references (*n* = 454, used to build `EGGMAN`’s Tier-2 profiles) reached 95.8% exact naming and 99.3% compendium concordance (zero wrong CRF calls; on `COMET`’s in-catalog `CRF01-118` subset, `COMET` scored 98.1% vs. `EGGMAN`’s 94.0%, while `EGGMAN`’s 81.8% match on 55 nominal pure references reflected reclassification of ancestral Subtype G/A7 mosaics and grouping of A3/A4/A8 within A1; Section 3.3).

Sharper differences emerged on out-of-sample clinical CRFs, leave-one-sequence-out (LOSO) cross-validation, and partial diagnostic reads. Across 150 out-of-sample clinical named CRFs, `EGGMAN` essentially tied `COMET` on exact legacy CRF naming (72.0% vs. 71.3%; 69.7% across 495 LOSO CRFs), but traded higher conservative `URF` fallback (27.3%, 92.7% compendium-consistent) for near-zero misassignment (1/150 wrong CRF call) while correctly flagging 91.0% of clinical `URF`s (*n* = 100). Crucially, evaluating all three tools symmetrically against independent published LANL CRF parental dictionaries (A-collapsed; excluding circular hotspot panels) showed that `EGGMAN` recovered the exact parental subtype set on 94.6% of clinical CRFs (80.5% from Tier-1 coordinates alone; mean Jaccard 0.976), outperforming both `COMET` (77.9%, Jaccard 0.891) and `jpHMM` (77.9%, Jaccard 0.919) while maintaining 97.3% mosaic-state recall (vs. 71.3% for `COMET` and 99.3% for `jpHMM`).

Because historical compendium boundaries carry annotation uncertainty—causing `EGGMAN` to differ from 483 legacy LANL bootscan midpoints by a median of 292 nt versus 89 nt for `jpHMM` (Section 3.3)—simulated mosaics with known ground truth provided the cleanest test of localization accuracy and specificity. Across 500 simulated GTR + Г_4_ mosaics (900 true junctions, processed in a fraction of a second), `EGGMAN` matched `jpHMM`’s nucleotide precision (9.0 nt vs. 10.0 nt median error; 97.1% vs. 89.3% within 100 nt) while cutting spurious junctions more than fivefold (7.9% vs. 41.8%) and lifting exact parental-set accuracy from 51.6% to 91.4% (Jaccard 0.975 vs. 0.725). Equally critical for high-volume screening, across 1,300 strictly non-recombinant null simulations spanning standard evolution, 3× rate acceleration, and APOBEC3G/3F hypermutation, Tier 1 produced 0.0% false recombinants and Tier 2 maintained 99.85% pure accuracy with zero false CRFs.

The most consequential separation appeared when tools confronted lineages absent from their catalog. On recently designated CRFs post-dating `COMET`’s static catalog (CRF119-157, *n* = 94), `EGGMAN` achieved 93.6% in-catalog exact naming and 96.8% exact independent parental sets (68.1% from Tier-1 coordinates alone), whereas `COMET` dropped to 0.0% exact naming (13.8% parent set, Jaccard 0.628) and `jpHMM` reached 79.8% parent set (Jaccard 0.927). Under strict matched out-of-dictionary masking (`CRF01-118` only, *n* = 152; alongside 94.1% `URF` flagging upon lineage omission across 495 LOSO CRFs), `EGGMAN` and `COMET` tied on overall non-legacy abstention (144/152 = 94.7% each: 141 `URF` + 3 `OutOfPanel` for `EGGMAN` vs. 144 `unassigned` for `COMET`, with 8/152 = 5.3% false legacy calls each). Here the two-tier distinction is operational: while `COMET` attached optional legacy CRF hints without breakpoint coordinates to 107/144 `unassigned` calls (75.7% total legacy-hinted) and `EGGMAN`’s 25-trunk epidemiological trace noted an ancestral founder CRF backbone on 66.4% (101/152), `EGGMAN` paired every abstention with a coordinate barcode recovering exact independent LANL parent sets on 68.1% of held-out CRF119-157 versus 13.8% for `COMET` (Jaccard 0.903 vs. 0.628).

### 3.2 One coordinate framework generalizes across viral architectures, separating benign naming granularity from consequential blind spots

Configuring the identical compiled engine solely through external `PathogenSpec` packs extended coordinate mapping and lineage certification across segmented orthomyxoviruses and coronaviruses (Table 2 and Fig. 2; full cohort ledger in Supplementary Table S10). For Influenza A H5Nx, we evaluated the eight-segment pack (*γ* (*x*) = 0 at segment boundaries, intra-segment crossovers disallowed) against published USDA `GenoFLU` genotype assignments across 1,100 labeled SRA H5N1 genomes (`GenoFLU` defines the labeled reference standard at 100%). Processing all 1,200 SRA genomes in a fraction of a second (nearly 10^4^ seq/s, more than an order of magnitude faster than 4-thread `GenoFLU`; Fig. 2A–B), `EGGMAN` matched `GenoFLU` on 97.8% (1,076/1,100) of labeled isolates across all 10 genotypes—including 100% concordance on A1, B3.2, and D1.3, 98.7%–99.2% on A3, B2.1, B3.13, and B3.6, and 93.8%–95.3% on D1.8, C2.1, and D1.1—with zero cross-genotype misclassifications. All 24 non-exact calls were conservative abstentions (23 `OutOfPanel`, 1 `URF`) triggered when an individual segment fell below the pack’s 90% identity floor. By contrast, a consequential gap appeared when testing uncatalogued reassortants. Withholding B3.13 and D1.1 from both tools (*n* = 650) left `GenoFLU` unable to classify any isolate (100% Not_assigned), whereas `EGGMAN` flagged 100% as novel mosaics and recovered the exact eight-segment donor formula on 98.9% (643/650). On the 100 USDA-`unassigned` SRA assemblies in the same cohort, `EGGMAN` resolved 98% into structured calls (52 certified genotypes, 25 `URF`s with eight-segment barcodes, and 21 `OutOfPanel`, with 92% multi-segment mosaics; 99% mosaic rate across the companion `unassigned` subset), and independent `IQ-TREE 2` segment trees verified all 16/16 donor placements across D1.8 and novel EA3[NAm_B313:PB2 | NAm_D1:PA+NP+NA] isolates (69%–100% UFBoot). Streaming 3,980 complete 2020–2026 BV-BRC H5Nx assemblies in under 10 s likewise separated 2,756 certified reassortants, 388 clonal founders, and 402 non-2.3.4.4b/LPAI `OutOfPanel` genomes from 420 `URF`s, organizing 352 of those `URF`s into 20 recurrent uncatalogued genotypes including European/EURL lineages absent from regional North American catalogs (Fig. 2B).

**Figure 2:**
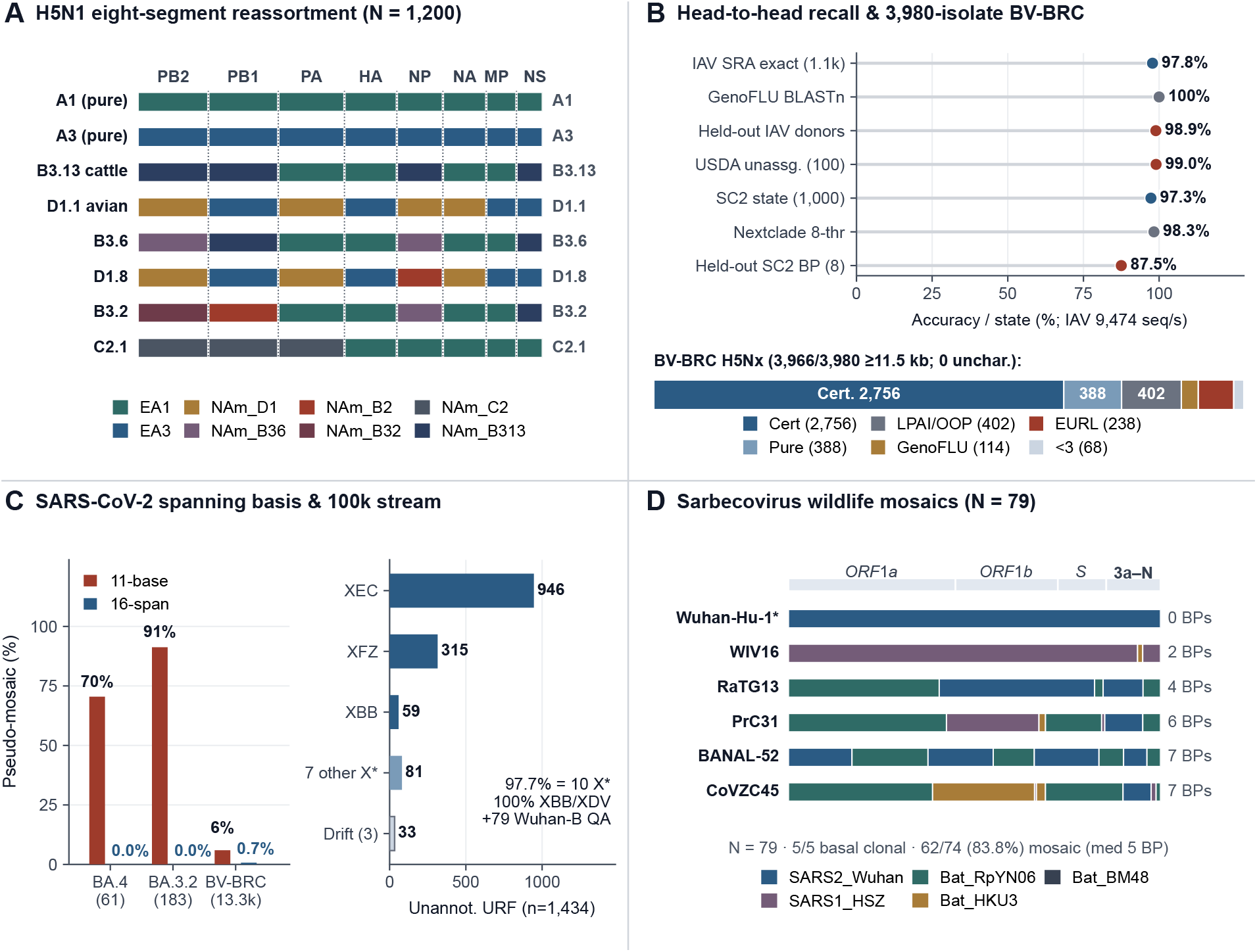
Cross-family recombination and reassortment profiling across Influenza A, SARS-CoV-2, and bat sarbecoviruses. **A:** Discrete *γ* (*x*) = 0 eight-segment mosaics across 1,200 SRA H5N1 genomes. **B:** Head-to-head accuracy vs. `GenoFLU` and `Nextclade` (top) and BV-BRC 3,980-isolate H5Nx partition into 2,756 certified, 388 pure, 402 `OutOfPanel`, and 420 `URF` (20 constellations + 68 rare) assemblies (bottom). **C:** Expanding SARS-CoV-2 from 11 to 16 spanning waves eliminates `BA.4` and `BA.3.2` pseudo-mosaics on the 10,062-genome basis-discovery union and cuts false flags eightfold across 13,275 (13.3k) nominal pure-wave genomes in the independent 23,376-genome BV-BRC stream (left), while CoV-tuned screening of 100,000 BV-BRC genomes recovers 97.7% (1,401/1,434) of clustered unannotated `URF`s across 10 Pango X* roots (right). **D:** Mosaic barcodes across 79 valid sarbecovirus genomes.

For SARS-CoV-2, we evaluated `EGGMAN` against curated Pango metadata labels across 1,000 clinical genomes (500 clonal waves and 500 named X* recombinants), using 8-thread `Nextclade` CLI as a specialist tree-based placer. Operating tree-free with just 28 reference profiles at order 10^3^ seq/s (roughly 3× faster single-threaded than 8-thread `Nextclade` including wrapper overhead), `EGGMAN` matched `Nextclade` on clonal-versus-recombinant discrimination (97.3% vs. 98.3%). Decomposing that performance across the two tiers shows both where classification is straightforward and how the two layers complement each other: on the 500 pure-wave genomes, `EGGMAN` achieved 99.4% (497/500) exact wave accuracy and kept 98.2% (491/500) breakpoint-free in Tier 1; on the 500 X* recombinants, Tier 1 localized *de novo* breakpoints directly in 81.0% (405/500, giving 89.6% combined Tier-1 clonal/mosaic accuracy), while Tier 2 rescued another 77 shallow-tract recombinants (principally XF and XBK) whose short terminal tracts fall below the *de novo* switch floor. Strict Tier-2 family concordance (80.4% vs. `Nextclade`’s 96.9%) reflected a single benign, conservative safeguard rather than clade confusion: 157 of `EGGMAN`’s 196 discrepancies were evolved XBB descendants routed to `URF` by Tier 2 even though 146/157 still received the exact XBB coordinate formula (BJ.1[BM.1.1.1:S..ORF10]) in Tier 1 (and 7 showed localized Wuhan-B Spike backfill). Conversely, tree-free coordinate mapping directly addressed three consequential surveillance failure modes. First, expanding Tier 1 from 11 to 16 waves under the Spanning Basis Rule (Fig. 2C) eliminated 70.5% (BA.4) and 91.3% (BA.3.2) pseudo-mosaics on the 10,062-genome global basis-discovery union and, on the independent 23,376-genome BV-BRC validation stream, cut nominal pure-wave false flags eightfold (5.96% → 0.72% CoV-tuned; 0.09% default across 13,275 pure-wave genomes). Second, coronavirus-tuned sensitivity (*γ* = 3.5, *m*_min_ = 3) detected *de novo* breakpoints in 7/8 held-out `XDV`/`XFZ`/XFC genomes and rescued 5/6 shallow four-SNP XEC recombinants. Third, streaming 100,000 independent live BV-BRC genomes (77,358 submitted in 2025–2026) in under two minutes unmasked 99%–100% of alias-hidden XBB recombinants, flagged localized post-2021 Wuhan-B amplicon-backfill artifacts (3 default; 79 CoV-tuned) via post-processing audit filters, and grouped unannotated `URF`s into 13 clusters (1,434 genomes), of which 10/13 (97.7% of clustered genomes, led by XEC, `XFZ`, and XBB) recovered 10 designated Pango X* recombinant roots without prior annotation.

Across deeper divergence in 79 valid bat and pangolin sarbecovirus genomes (completed in under a tenth of a second), all 5/5 basal reference trunks remained clonal while 83.8% (62/74) of non-basal wildlife genomes were classified as deeply mosaic (median 5.0 breakpoints), recovering published chimeric architectures across RaTG13, BANAL-52, PrC31, and WIV16 [10] (Fig. 2D). Finally, tree-free `ChronAeon` emergence dates closely tracked `IQ-TREE 2` phylogenies calibrated with `LSD2` and `TreeTime` (median absolute difference 0.33–0.42 yr across 32 SARS-CoV-2 clades—25 direct tree-free estimates plus 7 sparse clades using `LSD2` fallback, chronologically ordering 26/29 recombination DAG edges—and 0.16–0.18 yr across 15 Influenza A alignments; Table 2). In H5Nx, these emergence estimates established donor-versus-reassortant directionality on shared segments—placing Eurasian `EA1` (2021.9) and `EA3` (2020.1) ahead of B3.13 (2024.0) and D1.1 (2024.1), and dating NAm_B313 (2023.87) immediately before its 2023.90 first sampling. Enabling `--temporal-guard` removed 100% of future-clade leakage across all 16 collection-date and submission-date ablations (eliminating up to 186 donor violations in BV-BRC H5Nx and 36 in SRA H5N1).

### 3.3 Auditing HIV-1 baselines clarifies ancestral mosaics, emergence chronology, and a hub-dominated recombination genealogy

Screening the 2023 LANL HIV-1 Subtype Reference Alignment with reference-free `RhizAeon` and `EGGMAN` Tier-1 mapping separated benign sub-clade naming conventions from consequential ancestral mosaicism embedded inside nominal pure references (Fig. 3A; Supplementary Table S3). To avoid ad hoc editing, we applied an explicit scrubbing rule: tracts spanning conserved non-loop genes with reproducible inter-subtype ancestry outside hypervariable *gp120* loops were masked or demoted, whereas short or unindexed hits confined to hypervariable *gp120 V1–V5* loops (such as isolated loop signals in canonical B, C, F1, H, and A2 references, which are already masked during Viterbi segmentation) were retained. Under this criterion, while Subtypes A3, A4, and A8 simply clustered within A1 without crossing parental boundaries, `RhizAeon` identified 12 foreign or divergent 3’ tracts across 10 nominal pure references (*p* ≤ 1.83 × 10^−6^), and 7/55 (12.7%) pure references typed directly as Tier-1 mosaics: all four Subtype G references shared the ancestral G0[`A1:gp41..nef`] architecture [15], all three Subtype A7 references showed multi-parent mosaicism, and Subtype L (AF286236) harbored a 3’ `K:nef` tract. Conversely, no `?E` tracts appeared upstream of *vif* (`HXB2` 5097) across any reference or archive mosaic, establishing that the unsampled `CRF01_AE` parental lineage is strictly 3’-restricted [24, 25]. Separating the scrubbed basal trunk G0 placed Group M emergence dates on a consistent timeline (Fig. 3B): across 21 clades (3,594 dated genomes; using a two-regime spline for Subtype B), tree-free `ChronAeon` dates correlated strongly with `IQ-TREE 2` clocks calibrated with `LSD2` (*p* = 0.814*, p* = 7.04 × 10^−6^; median absolute difference 12.72 yr) and `TreeTime` (*p* = 0.791; median difference 11.23 yr). Point estimates placed Subtype C (1951.4) and `CRF02_AG` (1956.1, 95% CI 1938.2–1971.0) earlier than Subtype A1 (1959.7) and nominal Subtype G (1961.5, 1947.0–1973.9)—a 5.4-yr ordering that falls within overlapping confidence intervals and HIV-1 clock uncertainty, remaining directionally consistent with phylogenetic reconstructions [15]—followed by major regional waves from `CRF01_AE` (1974.7) and CRF07_BC (1988.2) to CRF63_02A6 (2009.6).

**Figure 3:**
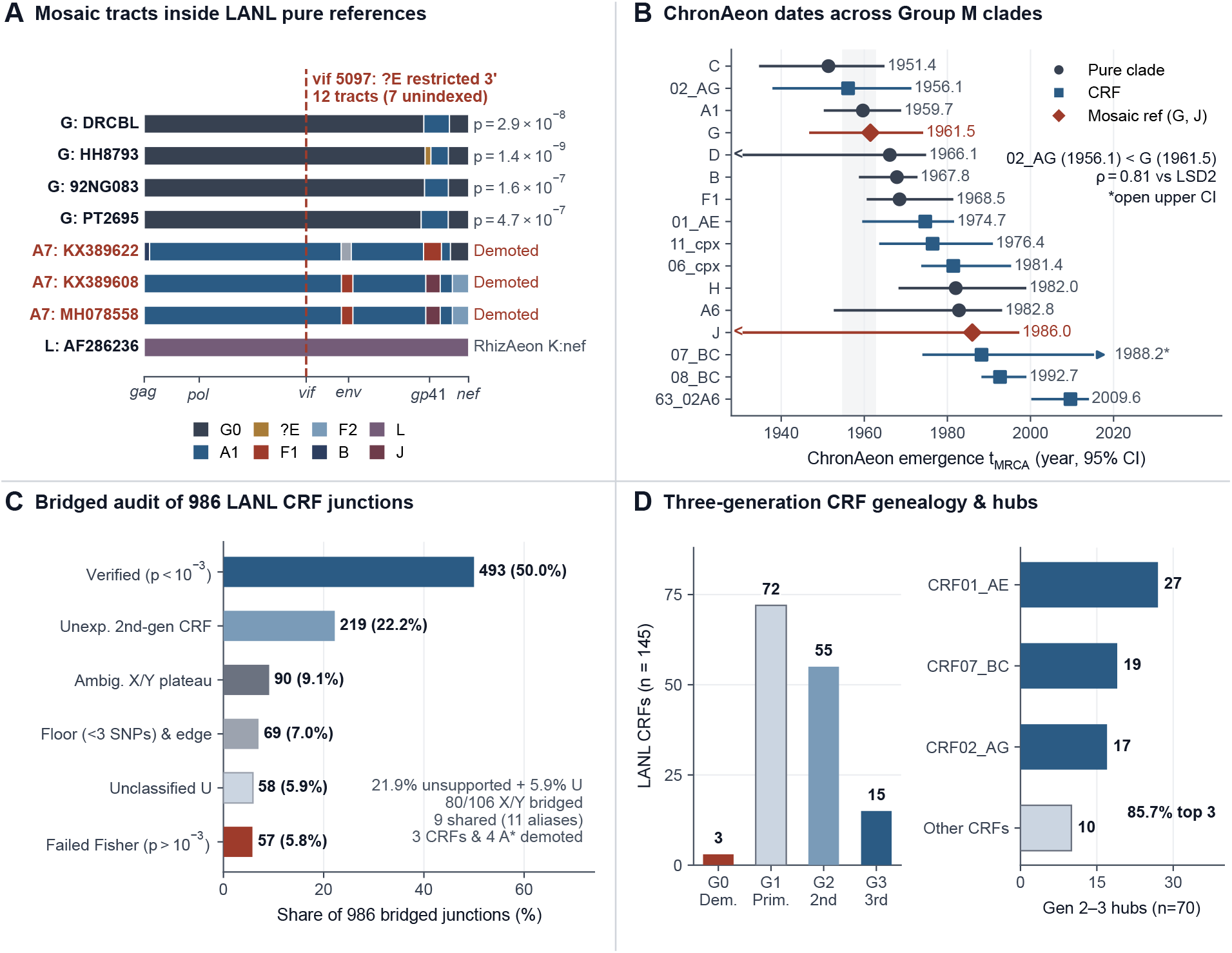
Harmonizing HIV-1 reference baselines, emergence chronology, CRF junctions, and three-generation hubs. **A:** Ancestral mosaic tracts inside canonical LANL pure references, including four Subtype G `G0[A1:gp41..nef]` genomes, three reclassified Subtype A7 chimeras, and strict `HXB2` 5097 restriction of `?E`. **B:** `ChronAeon` *t*_MRCA_ chronology across Group M clades (*p* = 0.814 vs. `LSD2`; median diff. 12.7 yr), placing `CRF02_AG` (1956.1) prior to Subtype G (1961.5) within overlapping CIs; left < markers on D and J denote lower CIs extending before 1931, and right arrows denote open upper bounds. **C:** Statistical audit of 986 testable LANL CRF junctions shows 493 (50.0%) verified direct switches, 219 (22.2%) junctions inherited from parent CRFs, 58 (5.9%) untestable unclassified U segments, and 216 (21.9%, or 23.3% of 928 indexed-donor junctions) statistically unsupported boundaries. **D:** Directed acyclic genealogy of 145 LANL CRFs reveals 3 retired/out-of-panel, 72 primary, 55 second-generation, and 15 third-generation CRFs, with 85.7% (60/70) of Generation 2–3 lineages tracing ancestry to `CRF01_AE` (27), CRF07_BC (19), and `CRF02_AG` (17).

Testing nearly a thousand published LANL CRF breakpoints (986 testable junctions; Fig. 3C and Supplementary Section S2) explained why modern coordinate segmentation differs from historical bootscan maps. Nearly three-quarters of catalogued junctions (72.2%) reflected genuine evolutionary boundaries—half (50.0%) representing direct crossovers between pure subtypes and another 22.2% inherited intact from an earlier parent CRF. Separating 58 (5.9%) junctions involving unclassified U donors—which are untestable against an incomplete Tier-1 basis rather than disproven switches—left 216 boundaries (21.9% of 986, or 23.3% of the 928 indexed-donor junctions) that lacked statistical support due to short or ambiguous intervals with too few informative substitutions. Translating sequential CRF numbers into coordinate formulas also revealed hidden structural overlap: nine distinct mosaic architectures are shared identically across 20 separately numbered CRFs, while three catalogued entries lacked statistically supported Tier-1 mosaics (CRF26_A5U and CRF92_C2U, involving unindexed parental lineages, and CRF46_BF1, which behaved clonally).

Ordering all 145 audited CRFs by parental descent into a directed acyclic genealogy revealed that global HIV-1 recombination is highly hierarchical rather than an even mixture of pure subtypes (Fig. 3D). Alongside 72 primary (Generation-1) CRFs formed directly from pure clades, nearly half of active CRFs (70/142, 49.3%) are second-generation (*n* = 55) or third-generation (*n* = 15) descendants built atop earlier CRFs. Remarkably, 85.7% (60/70) of these multi-generation lineages trace their ancestry back to just three major pandemic founder hubs: `CRF01_AE` (*n* = 27), CRF07_BC (*n* = 19), and `CRF02_AG` (*n* = 17).

### 3.4 Clinical and archive audits separate benign locus splits and convergent hotspots from genuine spreading lineages

Applying Tier-2 guarded subtyping to Stanford HIVDB [6] demonstrated how coordinate mapping separates benign locus-sampling effects from consequential classification discrepancies across 120,468 clinical *pol* records (Fig. 4A; Supplementary Section S3). On the 720-sequence reference panel, `EGGMAN` matched 91.7% of labels (98.3% of pure references, while flagging six nominal pure references carrying *pol* crossovers). Across the clinical surveillance archive, `EGGMAN` typed 8,563 previously unlinked submissions and agreed with 88.4% of 111,905 isolate-linked clinical entries. Auditing the remaining 11.6% (13,004) discrepancies separated expected diagnostic artifacts from genuine conflicts: nearly half (6,141, 5.5%) were benign multi-enzyme locus splits—where separate protease and reverse-transcriptase amplicons from the same mosaic virus straddle an internal CRF breakpoint (led by Chinese B vs. CRF07_BC, *n* = 2,396)—alongside 1,453 CRFs whose *pol* locus is non-mosaic and 1,243 ambiguous tags (principally Chinese B + C) clarified directly by coordinate calls. By contrast, only 2,157 records (1.9%) reflected true intra-patient clonal contradictions (led by Estonian CRF06_cpx vs. Subtype G, *n* = 520, and Ugandan B vs. D), alongside 1,444 nominal pure entries harboring genuine *pol* crossovers, 486 alternate CRF/trunk/01_AE assignments, and 80 (0.07%) short or unclassifiable fragments accounting for all 13,004 non-concordant linked entries. The audit also illustrated the practical value of Tier-2 stem certification on short clinical reads: across 218 West African `CRF02_AG` *pol* isolates whose bare Tier-1 trace drifted toward G0[A6:RT], Tier-2 guards assigned 87.2% (190/218) directly to `CRF02_AG`, corroborated by independent *pol.RT* maximum-likelihood phylogenies across 40/40 (100%) West African controls clustering with `CRF02_AG` rather than former-Soviet A6/CRF63 (0.069 vs. 0.156 substitutions/site).

**Figure 4:**
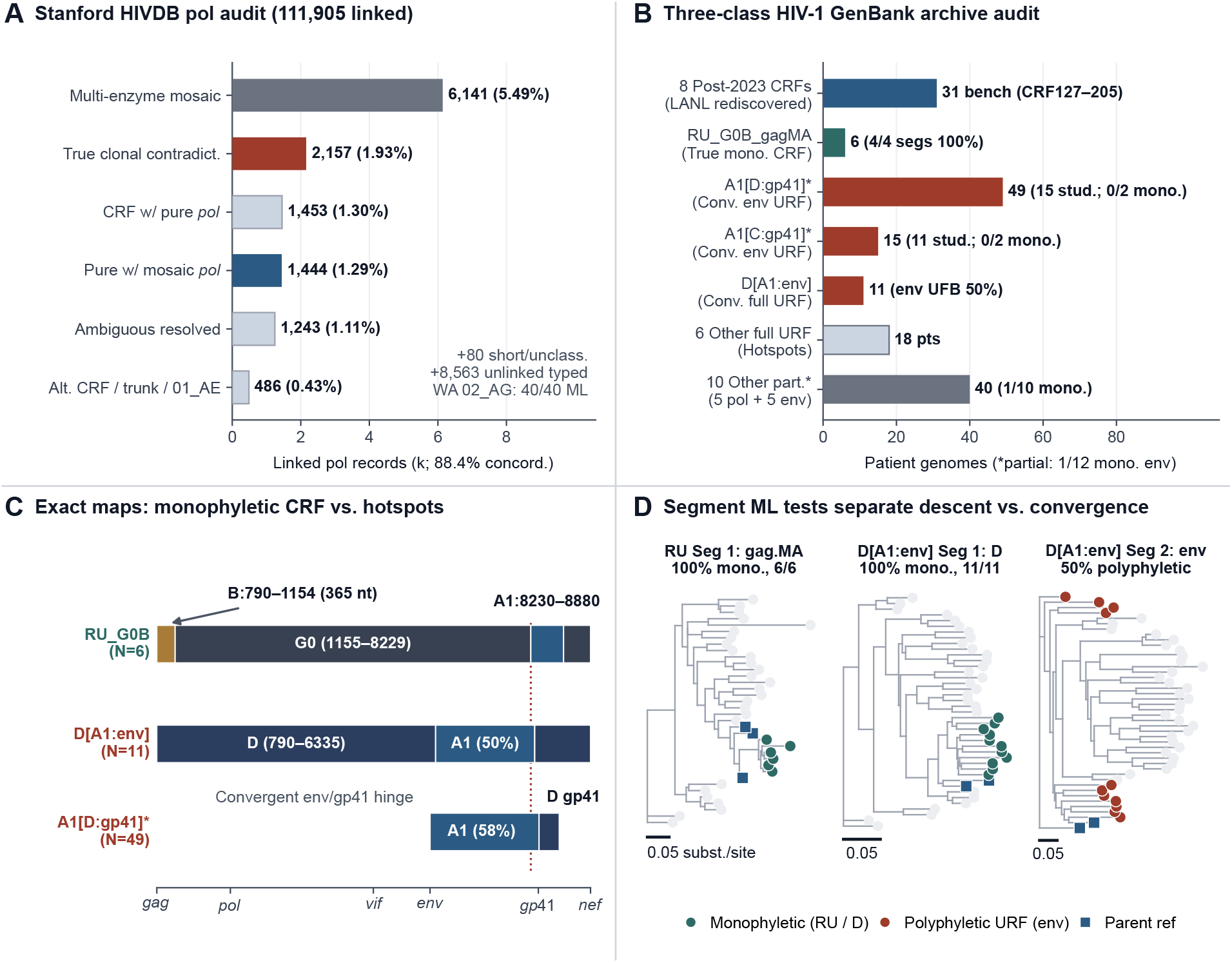
Stanford clinical *pol* reconciliation and three-class separation of true CRFs from convergent HIV-1 hotspots. **A:** Auditing 111,905 `isolates.d`-linked clinical *pol* entries (88.38% concordant; 13,004 non-concordant incl. 80 short/unclassifiable) separates expected multi-enzyme locus splits (6,141) from true intra-patient contradictions (2,157) and subtypes 8,563 unlinked records while validating 40/40 West African `CRF02_AG` ML trees. **B:** GenBank archive screening across 7,720 full-genome and 20,797 partial-only patients (7,897 *pol*, 12,900 *env*) distinguishes eight blindly rediscovered post-2023 LANL CRFs (31 benchmark genomes), one strictly monophyletic full-genome regional candidate (`RU_G0B_gagMA`, *N* = 6), one monophyletic partial-*env* cluster (`C[D:gp41]`, *N* = 3, Tanzania; full genome unavailable), and polyphyletic convergent hotspot `URF`s (7 full, 5 partial-*pol*, 6 partial-*env*). **C:** Exact true-scale `HXB2` maps contrast `RU_G0B_gagMA` (`B:gag.MA` 790–1154, 365 nt) against convergent D[`A1:env`] (*N* = 11) and partial `A1[D:gp41]` (*N* = 49, 15 studies) sharing the *env/gp41* junction. **D:** Segment ML trees confirm 100% monophyly across `RU_G0B_gagMA` vs. polyphyletic *env* convergence in `D[A1:env]` (50% UFBoot).

Scaling the same screen across NCBI GenBank (7,720 deduplicated full-genome hosts and 20,797 strictly partial-only *pol* or *env* hosts; Supplementary Tables S7–S8) showed that identical coordinate barcodes across unlinked patients do not automatically signify a single expanding epidemic (Fig. 4B–D and Supplemental Fig. S1). Screening all unindexed mosaic barcodes shared by at least three independent hosts (±100 nt breakpoints) and testing each cluster by segment-level `IQ-TREE 2` phylogenetics divided candidates into three distinct outcomes. First, eight full-genome clusters surfaced from pre-cutoff genomes matched official post-2023 LANL CRFs (CRF127_07109 through CRF205_0107; Fig. 4B), confirming prospective lineage detection. Second, beyond those official clades, exactly one unindexed full-genome regional cluster—six Russian genomes sampled during 2014–2016 (RU_G0B_gagMA, 4.49% mean divergence [35])—satisfied strict multi-segment monophyly and parental trunk concordance, sharing an exact three-parent architecture (G0[B:gag.MA | `A1:gp41..nef`], including a 365-nt B tract in *gag.MA*) supported at 100% UFBoot across all four segments (Fig. 4C–D and Supplemental Fig. S1A). Among 12 strictly partial-gene clusters (5 *pol*, 7 *env*), exactly one (1/12, Tanzanian C[D:gp41] across 3 patients from 3 independent studies) showed monophyly across both partial *env* segments (93% and 98% UFBoot), remaining unpromotable solely because full-genome sequence is absent (per Robertson et al. CRF criteria). Third, all remaining recurrent clusters—seven full-genome formulas and 11/12 partial-gene clusters (101/104 partial patients)—reflected repeated independent recombination at structural hinges (or divergent single-subtype sub-lineages, as in G0[H:gp41..nef]) rather than a shared recombinant transmission chain. Most prominently, East African full-genome D[A1:env] (11 patients across Uganda, Kenya, and Tanzania [36]) and 49 unlinked partial-*env* patients carrying A1[D:gp41] across 15 studies converged on the identical *gp120/gp41* hinge (median `HXB2` 8256), yet scattered polyphyletically across Subtype A1 in *env* trees (50% and 58% UFBoot; mirrored by A1[C:gp41] across 15 patients and 11 studies, 65% UFBoot; Fig. 4C–D and Supplemental Fig. S1B). Coordinate clustering thus surfaces recurrent mosaics rapidly at archive scale, while segment monophyly prevents mistaking convergent structural hotspots for circulating epidemics.

### 3.5 Census-level HIV-1 mapping reveals a two-stage selective sieve separating folding viability from epidemic transmission

Combining 7,720 deduplicated full-genome GenBank hosts with active CRFs turned archive-scale coordinate maps into a quantitative assay of recombinant emergence and survival (Fig. 5; complete statistical models and founder-deduplicated controls in Supplementary Table S9). Across 32 countries represented by at least 40 hosts, local circulating non-URF lineage diversity (*H*_circ_) strongly predicted *de novo* `URF` prevalence (Spearman *p* = 0.59*, p* = 3.9 × 10^−4^; Fig. 5A), remaining highly significant after adjusting for Sub-Saharan African geography, sample size, and sampling span across weighted least-squares (*R*^2^ = 0.62*, p* = 1.6×10^−4^), heteroskedasticity-consistent binomial (HC3, *p* = 1.5 × 10^−7^), and negative-binomial regression. Yet high co-infection supply rarely translates into sustained transmission: across 761 distinct mosaic lineages, observed host cluster sizes exhibited a steep heavy-tailed decline spanning three orders of magnitude (approximated by a discrete Hurwitz-zeta tail *α*^ = 2.21 ± 0.05, though statistically comparable to a discrete lognormal distribution and rejected as a strict single-exponent power law from *x*_min_ = 1 by Clauset bootstrap goodness-of-fit; Fig. 5B). Nearly three-quarters of mosaic lineages (71.6%, 545/761) remained single-host dead ends (491 `URF`s and 54 singleton CRFs), 19.1% (145) circulated in ≥ 3 hosts, and only 4.1% (31) reached ≥ 10 hosts—proportions that remained invariant when varying the host deduplication threshold nearly fourfold (0.4%, 0.8%, and 1.5% divergence; Supplementary Section S5).

**Figure 5:**
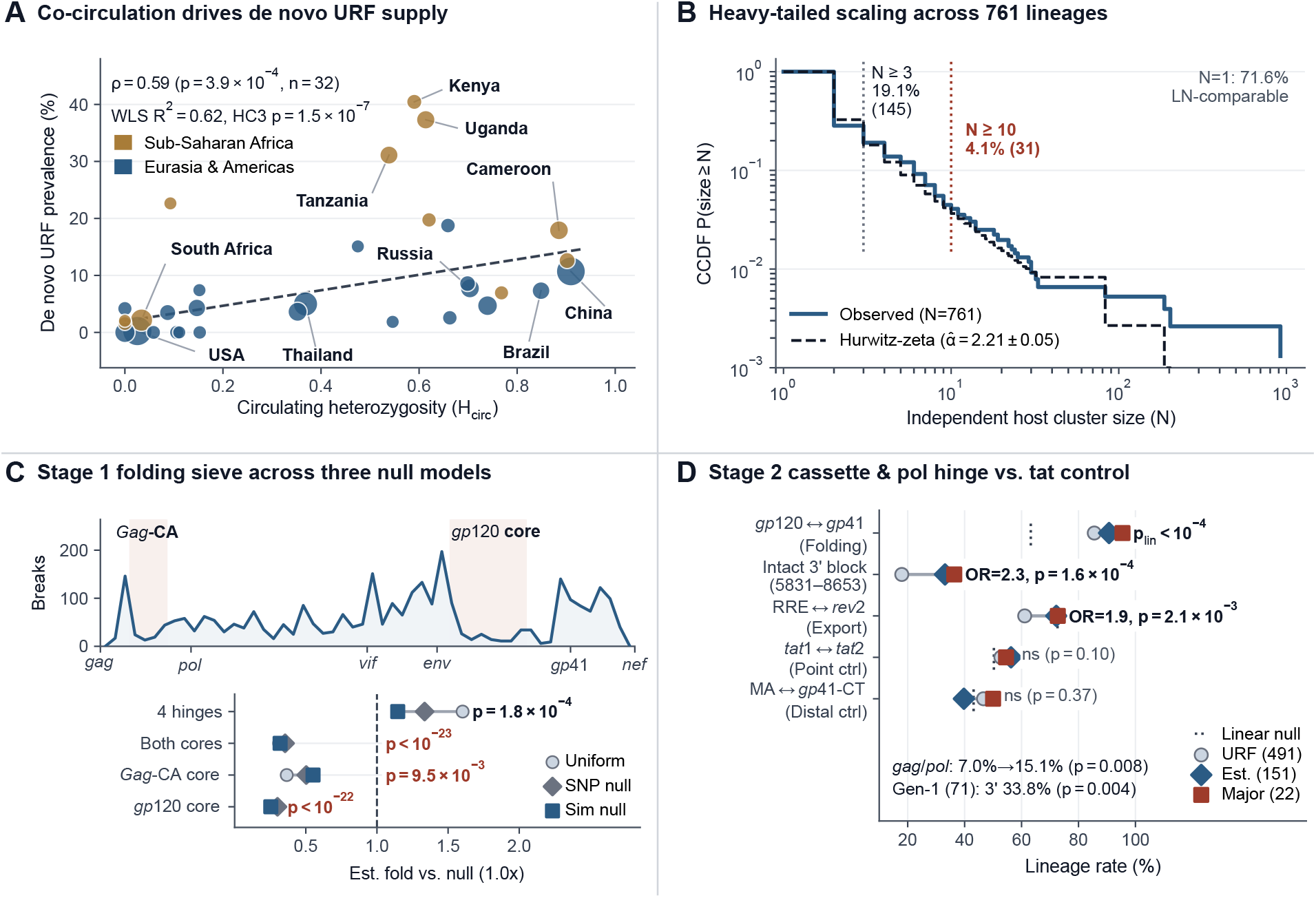
Co-infection supply, heavy-tailed cluster scaling, and a two-stage selective sieve across HIV-1 mosaics. **A:** National `URF` prevalence vs. non-URF heterozygosity *H*_circ_ across 32 countries (*p* = 0.589, WLS *R*^2^ = 0.621, HC3 GLM *p* = 1.45×10^−7^). **B:** Offspring cluster sizes (*N* = 761) show a steep heavy-tailed decline (Hurwitz-zeta *α*^ = 2.21 ± 0.05; comparable to discrete lognormal). **C:** Genome breakpoint flux (top) and Stage 1 fold depletion/enrichment relative to uniform, diagnostic-SNP, and `EGGMAN` detectability nulls (bottom) confirm avoidance of both *gp120* and *Gag*-CA cores. **D:** Across singletons (491), established (151), and major (22) CRFs, Stage 1 *gp120*–*gp41* folding lock holds across all tiers while Stage 2 transmission preserves an intact 3′ regulatory/Env cassette (5831–8653; all CRFs OR = 2.27, MV *p* = 9.6 × 10^−6^; Gen-1 founder-deduplicated *N* = 71: 33.8%, MV OR = 2.58*, p* = 0.004) and *RRE*–*rev2* linkage (Gen-1 81.7%, MV *p* = 0.004) rather than point *tat1*–*tat2* linkage (ns).

Comparing breakpoint locations across 491 singleton dead-end `URF`s (2,046 breakpoints), 151 established CRFs (701 breakpoints), and 22 major CRFs (*N*_host_ ≥ 10, 93 breakpoints) revealed a stringent Stage-1 protein-folding filter operating prior to onward transmission (Fig. 5C). Relative to uniform coding span, empirical diagnostic-SNP density, and `EGGMAN` detectability nulls (both 2%-noise single-crossover simulations and realistic 4-crossover simulations at 6%–10% GTR + Г_4_ divergence recalling 4,126/4,000 junctions), viable mosaics strongly avoided crossovers inside both the *gp120* ectodomain *β*-sandwich core (0.25–0.30× across nulls, *p* ≤ 1.6 × 10^−16^; 0.27× under 4-break 8%-divergence null) and the helical *Gag*-CA capsid core (0.36–0.55×, *p* ≤ 0.0095). This dual core depletion replicated under canonical full-gene boundaries (*gag.CA* 0.46×, *env.gp120* 0.63×; Supplementary Table S9) and appeared in consistently high co-ancestry between *gp120* and the *gp41* ectodomain across every survival tier (85.5% in `URF`s rising to 95.5% in major CRFs, *p*_lin_ *<* 10^−4^), whereas four flexible inter-domain hinges absorbed 55.8%–60.8% of surviving crossovers (1.15 × –1.47× vs. detectability nulls, *p* ≤ 1.8 × 10^−4^), consistent with structural folding-preservation constraints [37, 38]. By contrast, the *Pol*-RT palm/fingers region (2650–3450) proved crossover-tolerant rather than depleted in established CRFs (11.7% overall, 12.0% across 527 DAG-deduplicated *de novo* CRF breakpoints vs. 6.7% in `URF`s, *p* = 1.2 × 10^−4^).

Contrasting established and major CRFs directly against singleton `URF`s then isolated the Stage-2 filter governing between-host transmission—explicitly controlling for phylogenetic pseudoreplication from Generation 2–3 CRF hubs by re-evaluating every signal across the 71 alias-collapsed Generation-1 primary founder CRFs alongside parental-pair–stratified CMH and multivariable logistic models adjusting for break-point count and parental trunk pair (Fig. 5D; Supplementary Table S9). Progression to an established CRF was not driven by isolated two-point co-ancestry between *tat* exon 1 and *tat* exon 2 (52.7% in `URF`s vs. 56.3% in all established and 63.4% in Gen-1 primary CRFs; all-CRF CMH *p* = 0.42, MV *p* = 0.16; Gen-1 CMH *p* = 0.16, MV *p* = 0.093). Instead, the clearest survival signal surviving both founder deduplication and multivariable adjustment was preservation of the entire 3’ spliced regulatory and envelope cassette (`HXB2` 5831–8653: *tat1–vpu–env–tat2–rev2*) completely undivided—nearly doubling from 17.9% (88/491) in dead-end `URF`s to 33.1% (50/151) in all established CRFs (CMH OR = 2.20*, p* = 1.2 × 10^−3^; multivariable OR = 3.09*, p* = 9.6 × 10^−6^), 36.4% (8/22) in major CRFs, and 33.8% (24/71) in strictly independent Gen-1 primary CRFs (CMH OR = 2.11*, p* = 0.025; multivariable OR = 2.58*, p* = 4.0 × 10^−3^). Within that 3’ cassette, *RRE*–*rev2* co-ancestry strengthened further upon removing multi-generation hub descendants—rising from 61.1% in `URF`s to 81.7% (58/71) in Gen-1 primary CRFs (exceeding its own 73.6% Gen-1 linear permutation null, *p* = 0.046; CMH OR = 2.31*, p* = 0.021; multivariable OR = 2.67*, p* = 4.0 × 10^−3^; 72.2% across all 151 CRFs, MV *p* = 0.027 adjusting for breakpoint count and pair, attenuating to *p* = 0.119 when conditioning on colinear 8150–8550 breaks). By contrast, founder-deduplicated and stratified controls showed that two secondary univariate patterns were largely confounded by parental ancestry and hub inheritance: crude suppression of *gp41*-CT return swaps (35.6% in `URF`s vs. 19.5% in all CRFs, *p* = 0.0042) vanished under parental-pair stratification (CMH OR = 0.79*, p* = 0.60; Gen-1 CMH OR = 0.86*, p* = 0.97), while upstream *gag/pol* enrichment (7.0% → 10.7% → 15.1% major, all-CRF CMH *p* = 0.008; 9.7% across DAG-deduplicated *de novo* junctions, *p* = 0.042) attenuated in Gen-1 primary founders alone (9.3%*, p* = 0.16). Together, these founder-independent controls pinpoint preservation of the intact 3′ transcriptional, nuclear-export, and envelope module as the primary reproducible Stage-2 correlate of epidemic persistence [39–42].

## 4 Discussion

Treating viral sequence classification as a single flat naming problem conflates two distinct evolutionary tasks: reconstructing parental ancestry along physical genome coordinates (Tier 1) and testing whether an isolate descends from an established epidemic clone (Tier 2). Separating these operations behind external `PathogenSpec` packs allows `EGGMAN` to complement existing community frameworks rather than replace their vocabularies. Familiar clinical and epidemiological designations (`CRF02_AG`, B3.13, XEC) are preserved whenever a sequence clears four sequential statistical guards, while newly emerging or partial-genome mosaics immediately receive transparent, Day-1 coordinate identifiers (`Backbone[Donor:Region]`) at 10^3^–10^4^ sequences per second natively or in a browser. Our cross-pathogen benchmarks also clarify how different classes of tools divide labor effectively: full phylogenetic placers such as `Nextclade` remain well suited for fine-grained sublineage naming within a dense single-pathogen tree, whereas `EGGMAN` provides a fast, portable coordinate layer for *de novo* breakpoint deconvolution, alias unmasking, sequencing-artifact screening, and multi-segment reassortment tracking.

Auditing public repositories at whole-archive scale underscores two general principles for genomic surveillance. First, reference panels benefit directly from systematic recombination and temporal hygiene: masking ancestral chimeric tracts such as `A1:gp41..nef` inside Subtype G restores consistent Group M emergence chronologies, applying the Spanning Basis Rule prevents newly diverged pure waves (BA.4, BA.3.2) from inflating pseudo-mosaic rates, and opt-in temporal guarding prevents retrospective lookahead bias in longitudinal evaluations. Second, fast coordinate clustering functions best as an archive-scale discovery filter paired with phylogenetic validation rather than a standalone criterion for naming new lineages. Across more than 28,500 HIV-1 hosts in GenBank, shared unindexed coordinate barcodes surfaced both genuine circulating clades—eight post-2023 official LANL CRFs and the strictly monophyletic Russian cluster RU_G0B_gagMA—and polyphyletic convergent hotspots such as East African D[A1:env] and A1[D:gp41]. Coupling coordinate maps with multi-segment monophyly checks prevents mistaking recurrent recombination hotspots for single expanding transmission chains.

At the same time, uniform coordinate mapping turns public sequence archives into a quantitative assay of viral evolution. Across 761 HIV-1 mosaic lineages, local circulating subtype diversity governs the ecological supply of *de novo* recombinants, yet severe transmission bottlenecks restrict more than 70% of newly formed mosaics to single hosts. Controlling explicitly for sequence divergence, algorithmic detectability, and Generation 2–3 CRF hub pseudoreplication demonstrates that survival passes through two distinct selective filters: within hosts, structural folding constraints purge crossovers inside both the *gp120 β*-sandwich and the helical *Gag*-CA capsid core while concentrating viable switches inside flexible inter-domain linkers [37, 38]; between hosts, epidemic persistence acts not through an isolated two-exon *tat1*–*tat2* lock, but primarily by preserving the integrated 3′ transcriptional, nuclear-export, and envelope module (*tat1–vpu–env–tat2–rev2*, including *RRE*–*rev2* linkage in primary founders) undivided [39–42].

Several practical boundaries define how `EGGMAN` should be applied and extended. First, Tier-1 coordinate reconstruction depends on a representative non-mosaic founder basis: when substantially novel out-of-panel clades emerge, conservative `Unattributed` and `URF` flags appropriately signal novelty rather than fabricating ancestry, requiring periodic pack updates governed by the Spanning Basis Rule as viral populations diversify. Second, discrete Viterbi segmentation trades site-level posterior uncertainty for multi-order-of-magnitude speed; for low-divergence recombinants carrying short terminal tracts (such as shallow SARS-CoV-2 XF, XBK, or XEC mosaics), sensitivity depends on pathogen-tuned switch penalties or Tier-2 stem profiles, and fine-scale single-isolate breakpoint boundaries still benefit from follow-up phylogenetic or probabilistic analysis (`GARD`, RDP5, `RhizAeon`). Third, because census-level HIV-1 analyses rely on public repositories with uneven geographic sampling, targeted longitudinal cohorts will help refine estimates of between-host recombinant transmissibility. Looking forward, decoupling the engine from external `PathogenSpec` packs allows domain communities to maintain versioned, emergence-dated specifications across additional recombining and segmented viruses—from enteroviruses and flaviviruses to seasonal influenza and emerging zoonotic coronaviruses—supporting a consistent, reproducible coordinate layer for genomic epidemiology.

## Data and Code Availability

Source code for the `EGGMAN` Rust CLI and WebAssembly browser library (v0.2.0), audited HIV-1 reference profiles (152 active Tier-2 callers among 171 catalogued entries), external `PathogenSpec` packs (packs/ influenza_a_8seg, packs/sars_cov2_wuhan, packs/sarbecovirus_wuhan), and reproducible analysis scripts are publicly available at https://github.com/veg/eggman and https://veg.github.io/eggman/. The complete, versioned replication package—containing pinned software and packs, all machine-readable supplementary data tables (Data_S1–Data_S7: full `ChronAeon` vs. `LSD2`/`TreeTime` dates across 21 HIV-1, 32 SARS-CoV-2, and 15 Influenza A alignments; the 145-CRF genealogy DAG; the 32-country *H*_circ_ and GLM table; Generation-1 founder-deduplicated sieve and deduplication-sensitivity ledgers; non-circular benchmark outputs; 131 `IQ-TREE 2` segment phylogenies; and public accession manifests with archive retrieval dates), redistributable synthetic and HIV-1 benchmark cohorts, and a single-command verification harness—is archived on Zenodo at https://doi.org/10.5281/zenodo.24405743.

## Acknowledgments

We thank genomic surveillance contributors across LANL, NCBI GenBank, Sequence Read Archive, BV-BRC, and the Stanford HIV Drug Resistance Database whose public sequence archives enabled multi-pathogen coordinate indexing.

## Author contributions

S.L.K.P. and D.P.M. conceived the study and designed the coordinate nomen-clature framework. S.L.K.P., S.W., D.C., J.D.Z., and A.G.L. implemented `EGGMAN` and the interactive web workbench. S.L.K.P., D.C., J.D.Z., A.G.L., A.N., and D.P.M. curated multi-pathogen reference panels and conducted empirical benchmarks and evolutionary analyses. S.L.K.P., A.N., and D.P.M. wrote and revised the manuscript with input from all authors.

## Use of generative artificial intelligence

During the preparation of this work, the authors used large language model coding and writing assistants (Google Gemini via Antigravity) under direct author supervision to assist with software engineering, benchmark automation, figure layout scripting, LaTeX formatting, and prose editing. No generative AI tool is listed as an author, and no synthetic or AI-generated data were used in any empirical benchmark; all scientific hypotheses, algorithmic specifications, empirical analyses, statistical inferences, and manuscript claims were independently audited, executed on authentic public sequence datasets, and verified by the human authors, who accept full responsibility for the integrity and accuracy of the work.

