## Supplementary Material for "Come together: Coordinate deconvolution of viral recombination, reassortment, and epistatic survival"


This PDF file includes:

- Supplementary Materials and Methods
- Supplementary Text (Sections S1 to S6)
- Supplementary Figure S1
- Tables S1 to S10
- References

**Zenodo Replication Package (https://doi.org/10.5281/zenodo.24405743):** Machine-readable supplementary data tables (`Data_S1`–`Data_S7`, covering all `ChronAeon` vs. `LSD2`/`TreeTime` chronologies, the 145-CRF genealogy DAG, 32-country $H_{\text{circ}}$/GLM covariates, Generation-1 founder-deduplicated sieve and deduplication-sensitivity ledgers, non-circular multi-pathogen benchmark tables, Stanford HIVDB and GenBank cluster audits, and public accession lists with retrieval dates), 131 `IQ-TREE 2` segment trees, pinned `EGGMAN v0.2.0` source and `PathogenSpec` packs, and single-command verification scripts are archived permanently on Zenodo.

#### Supplementary Materials and Methods

##### S1.1 Sequence Curation and Multi-Pathogen `PathogenSpec` Hierarchy

Public HIV-1 sequence curation spanned four independent repositories: the 2023 Los Alamos National Laboratory (LANL) Subtype Reference Alignment ($N$ = 518), the LANL filtered full-genome alignment

($N$ = 5,088), NCBI GenBank full-length HIV-1 records ($N$ = 31,825, collapsing < 0.8% Hamming duplicates within identical mosaic barcodes to 7,720 independent patients), strictly partial-only GenBank and LANL *pol* (10,365 sequences = 7,980 GenBank + 2,385 LANL across 7,897 deduplicated hosts) and *env* (27,885 sequences = 23,163 GenBank + 4,722 LANL across 12,900 deduplicated hosts; 38,250 sequences and 20,797 partial-only patients total), and the Stanford HIV Drug Resistance Database (HIVDB) [1], comprising the reference panel (`genotype-references.fasta`, $N$ = 720) and clinical surveillance archive (`hiv-survdb`, $N$ = 120,468 partitioned into 111,905 `isolates.d`-linked entries and 8,563 [7.11%] unlinked FASTA submissions).

Cross-family validation configured external `PathogenSpec` reference panels (`--pack`) without modifying segmentation code:

1. *Influenza A H5Nx (`influenza_a_8seg`, $L$ = 13,136 nt across PB2–NS):* 10 `ChronAeon` time-decay donor trunks ($t_{1/2}$ = 0.50 yr; four Eurasian `EA1–EA4` and six North American `NAm_*`) paired with 10 Tier-2 profiles (2 pure `A1/A3` + 8 reassortant genotypes: `B3.13`, `D1.1`, `D1.3`, `B3.6`, `C2.1`, `B2.1`, `D1.8`, `B3.2`), evaluated across 1,200 complete SRA field isolates (1,100 USDA `GenoFLU` [2]-labeled + 100 "Not assigned" assemblies), 650 held-out `B3.13/D1.1` OOD tests, and 3,980 complete eight-segment BV-BRC 2020–2026 assemblies (34,384 segment rows assembled; 3,966 high-coverage ≥ 11,500 nt).

2. *SARS-CoV-2 (`sars_cov2_wuhan`, $L$ = 29,903 nt Wuhan-Hu-1 coordinates; coding 266–29674):* 16 `ChronAeon` time-decay Tier-1 spanning founders ($t_{1/2}$ = 0.25 yr) paired with 28 circulating Tier-2 profiles (12 pure waves + 16 recombinants; `BJ.1`, `BM.1.1.1`, `KP.3.3`, and `KS.1.1` serve exclusively as Tier-1 recombinant parents), evaluated across 1,000 curated clinical genomes (head-to-head against 8-thread `Nextclade CLI` [3]), 8 held-out `XDV/XFZ/XFC` OOD tests, 10,062 Global World Union genomes, and two live BV-BRC streams (23,376 stratified and 100,000 live genomes containing 77,358 2025–2026 entries).

3. *Wildlife Sarbecoviruses (`sarbecovirus_wuhan`, $L$ = 29,904 nt):* 5 basal non-recombinant trunks evaluated over 83 published whole genomes (79 valid genomes after rejecting four all-`N` GISAID placeholders `EPI_ISL_852604`, `852605`, `412977`, and `410721`) [4].

#### S1.2 Reference Scrubbing (`RhizAeon`), `ChronAeon` Dating, Temporal Guards, and Spanning Basis Selection

Because `RhizAeon` [5] and `ChronAeon` [6] are companion methods under preparation, we summarize both algorithms self-containedly here. `RhizAeon` screens a multiple alignment of candidate founder clades without an external reference tree by computing per-window pairwise and clade-consensus distance incongruence profiles, locating candidate donor shifts via dynamic programming, and testing each proposed boundary with an upper-tail $2 \times 2$ Fisher exact test on informative flanking substitutions. To prevent ad hoc curation, we applied one objective scrubbing rule: tracts spanning conserved non-loop genes with reproducible inter-subtype ancestry outside hypervariable *gp120* loops were masked or demoted (yielding `G0` with `A1:gp41..nef` masked, Subtype L with `K:nef` masked, demoted Subtype A7, and `?E` restricted to HXB2 5097–9417), whereas short or unindexed hits confined to hypervariable *gp120 V1–V5* loops (already masked during Viterbi scoring) were retained.

`ChronAeon` estimates clade root profiles and emergence times ($t_{\mathrm{MRCA}}$) directly from dated alignments without phylogenetic tree search. Each dated sequence $i$ (sampling time $t_i$, earliest sample $t_0$) receives an exponential time-decay weight $w_i = 2^{-(t_i - t_0)/t_{1/2}}$ ($t_{1/2}$ = 6.93 yr HIV-1, 0.50 yr IAV, 0.25 yr SARS-CoV-2), yielding deletion-aware base frequencies $Q(x, b)$ (assigning `N` where weighted gaps exceed valid bases) and per-sequence expected mismatch $P_{iQ}$ across valid sites $\mathcal{V}_i$ ($|\mathcal{V}_i| = L_i$):

$$Q(x,b) = \frac{\sum_i w_i \, \mathbb{I}[b_i(x) = b]}{\sum_i w_i \, \mathbb{I}[b_i(x) \in \{\mathrm{A,C,G,T}\}]}, \qquad P_{iQ} = \frac{1}{L_i} \sum_{x \in \mathcal{V}_i} \big[1 - Q(x, b_i(x))\big]. \tag{1}$$

Defining the mean site heterozygosity of the root profile as $h_{QQ} = \frac{1}{L}\sum_{x=1}^{L}\big[1 - \sum_{b\in\{\mathrm{A,C,G,T}\}} Q(x,b)^2\big]$, subtracting standing ancestral polymorphism $P_{rQ} = \frac{1}{2}h_{QQ}$ yields the adjusted root-to-tip mismatch proportion $P_{ri} = \max(0,\ P_{iQ} - P_{rQ})$, which converts to Jukes–Cantor Gamma distance ($\alpha_\Gamma = 0.20$):

$$d_\Gamma(P_{ri}) = \frac{3}{4}\alpha_\Gamma \left[\max\!\left(0.05,\ 1 - \frac{4}{3}P_{ri}\right)^{-1/\alpha_\Gamma} - 1\right], \tag{2}$$

followed by OLS regression of $d_\Gamma(P_{ri})$ on $t_i$ (using a two-regime spline for HIV-1 Subtype B and `LSD2_Fallback` for 7/32 sparse SARS-CoV-2 clades lacking sufficient temporal signal), 500–2,000 block bootstraps, and Fieller CIs [7]. Under `--temporal-guard` with `--temporal-mode sampled|submission`, a Tier-1 donor or Tier-2 catalog clade $k$ is admitted for a query dated $t_q$ iff $t_{\mathrm{floor}}(k) \le t_q + \delta_t$. In `sampled` mode, $t_{\mathrm{floor}}(k)$ uses earliest collection date (falling back to $t_{\mathrm{MRCA}}$) with pack slack $\delta_t = 3.0$ yr (HIV-1) or 0.5 yr (non-HIV); in `submission` mode, $t_{\mathrm{floor}}(k)$ uses earliest public deposition date with $\delta_t = 0.0$ yr (falling back to earliest sampled date on HIV-1 Tier-1 basal trunks).

For rapidly evolving outbreaks, the Spanning Basis Selection Rule evaluates each candidate clonal clade $c$ on $\le 25$ early representatives against existing Tier-1 basis $\mathcal{B}$, promoting $c$ into Tier 1 iff: (a) *separability*, $\min_{b\in\mathcal{B}} d_{\mathrm{coding}}(c,b) \ge 2m_{\min}$; (b) *irreducibility*, screening $c$ over $\mathcal{B}$ yields no statistically verified breakpoint; and (c) *completeness*, $> 50\%$ of early $c$ genomes would otherwise receive pseudo-mosaic calls or $c$ carries $\ge 25$ private coding substitutions relative to $\mathcal{B}$.

#### S1.3 Seed-Chained Bitmask Alignment, Split-DP, Unit-Mismatch Viterbi, and Breakpoint Support

Parameters in `EGGMAN` are chosen from sequence combinatorics and empirical viral substitution rates without fitting against benchmark test sets (Table S1). Exact 15-mer seeds (step 2 nt across indexed reference profiles, yielding random collision probability $4^{-15} = 9.31 \times 10^{-10} \ll 1/L$) are chained via longest increasing subsequence on both forward and reverse-complement strands, selecting the higher-scoring orientation (`align_unaligned_to_ref`; auto-pack detection in WASM uses a $< 15\%$ forward-hit fast path in `kmer_hit_stats`). When $< 4$ 15-mer hits occur ($q_{\mathrm{len}} \ge 80$), an 11-mer fallback pass (profile step 3 nt, query step 2 nt) runs, supplemented by local step-1 11-mer rescue across unanchored gaps or tails $> 350$ nt. Between consecutive anchors $\{(i_m, x_m)\}$, banded Needleman–Wunsch operates within diagonal radius $B = 45$ nt (50 nt on free tails):

$$|(x - x_m) - (i - i_m)| \le B, \qquad B = 45 \text{ nt}, \tag{3}$$

scoring exact `A/C/G/T` matches against the 4-bit reference-profile union mask (`base_mask_ref`) at +2, mismatches at −1, and linear gaps at −3 per nucleotide (query IUPAC ambiguity symbols decode to −1 and are skipped with 0.0 cost during Viterbi ancestry scoring). Whenever $|(x_{m+1} - x_m) - (i_{m+1} - i_m)| > 180$ nt, a forward/backward unit-diagonal split-DP solver (`STRUCT_BAND = 45 nt`) maximizes $F(p) + R(p)$ over the split coordinate $p$ to jump cleanly across large insertions or deletions without gap-smearing.

Across the projected coding span ($L$ = 8,628 nt, HXB2 790–9417 for HIV-1), discrete Viterbi dynamic programming assigns each coordinate $x$ a hidden state $k \in \{1, \ldots, K, \mathrm{Unattr}\}$ (for HIV-1 Tier 1, $K = 14$: 13 scrubbed pure Group M trunks plus locus-restricted `?E`):

$$V_k(x) = e_k(x, b_q(x)) + \min_{j\in\{1,\ldots,K,\mathrm{Unattr}\}} \Big[V_j(x-1) + \gamma(x)\,\mathbb{I}[j \ne k]\Big], \tag{4}$$

where costs are expressed in mismatch-equivalent (Hamming) units rather than log-likelihood nats: trunk emissions assign match cost 0.0, unambiguous mismatch cost 1.0, ambiguous/reference-gap cost 0.35, and out-of-range cost 2.0; `?E` carries prior offset $\delta_{?\mathrm{E}} = 0.035$ /site on HXB2 $x \geq 5097$ while being masked upstream; and `Unattributed` emits any query base at constant cost 0.58 /site (functioning as a high-divergence / alignment-failure floor above 58% mismatch). Under a symmetric per-site substitution probability $p$, a single mismatch corresponds to a log-likelihood ratio of $\ln[(1-p)/(p/3)]$ nats ($\approx 3.5$ nats at HIV-1 $p \approx 0.08$, versus $\approx 8.0$ nats at SARS-CoV-2 $p \approx 10^{-3}$); consequently, a fixed mismatch-unit switch penalty imposes a steeper probabilistic barrier in low-divergence viruses, directly explaining why contiguous presets scale with pathogen divergence ($\gamma = 7.0$ default HIV-1; 10.5 compendium auditing; 13.5 full-genome surveillance; 3.5 CoV-tuned). On segmented packs, $\gamma(x) = 0.0$ across segment boundaries and $10^6$ within segments when intra-segment recombination is forbidden. Every candidate junction between states $A$ and $B$ must satisfy flanking spans ($\geq 120$ nt full / 150 nt partial; $\min(L_{\min}, \lfloor L_{\mathrm{seg}}/2 \rfloor)$ segmented), $\geq m_{\min}$ informative SNPs (5 default; 3 CoV-tuned; 7 partial/full screen), and an upper-tail $2 \times 2$ Fisher exact test on left/right diagnostic counts $(n_{L,A}, n_{L,B};\, n_{R,A}, n_{R,B})$:

$$p_{\mathrm{Fisher}} = \sum_{u=n_{L,A}}^{\min(n_L, n_A)} \frac{\binom{n_L}{u}\binom{n_R}{n_A-u}}{\binom{n_L+n_R}{n_A}} < p_{\mathrm{thresh}} \qquad (10^{-4} \text{ default; } 10^{-3} \text{ comp.; } 0.01 \text{ CoV}). \tag{5}$$

#### S1.4 Four-Gate Lineage Certification, `OutOfPanel` Guards, and 25-Trunk Trace

Tier 2 evaluates 152 active HIV-1 callers (142 active 2023 LANL CRFs with `CRF31_BC`, `CRF35_A1D`, and `CRF146_BC` restored + 8 post-2023 CRFs `CRF127_07109`, `CRF135_0107`, `CRF136_0107`, `CRF162_cpx`, `CRF172_0755`, `CRF173_63A6`, `CRF188_0107`, `CRF205_0107` + `Subtype_G` + candidate `RU_G0B_gagMA`) within 171 catalogued entries (12 pure subtypes and 7 demoted entries: `A3`, `A4`, `A7`, `A8`, out-of-panel `CRF26_A5U`/`CRF92_C2U`, and clonal `CRF46_BF1`). Candidate lineage $c$ must clear four sequential checks:

1. *Gate 1 (Footprint & donor plurality):* $\geq$ 100 nt mapped at $\geq$ 84% sequence identity covering $\geq$ 2 donor blocks ($\geq$ 120 nt each when composite) with strict donor plurality $c_w > b_w$ on $\geq$ 3 informative SNPs per donor block.

2. *Gate 2 (Parental & DAG dominance):* against every pure constituent parent and immediate DAG ancestor $p$, net support $\Delta_2 = n_c - n_p \geq 5$ full (3 partial) and one-sided binomial probability $p_2 = \sum_{k=n_c}^{n} \binom{n}{k} 2^{-n} \leq 10^{-3}$ full ($10^{-2}$ partial; $n = n_c + n_p$).

3. *Gate 3 (Clonal stem excess):* contrasting $c$ against composite parental expectation $B_c(x)$ along $c$'s canonical barcode, net post-recombination stem excess $\Delta_{\mathrm{stem}} = s_c - s_B \geq s_{\min}$ (8 HIV full / 5 HIV partial / 4 IAV full [3 partial] / 2 SC2 full [1 partial]) with one-sided binomial $p_3 \leq 10^{-4}$ full ($10^{-2}$ partial; 0.05 IAV/SC2).

4. *Gate 4 (All-excursion Kadane residual scan):* for each competing clade $k$ over residual indicators $r_x \in \{+1, -1, 0\}$, evaluating $S_{\max}(k) = \max_{i \leq j} \sum_{x=i}^{j} r_x$ and rejecting any $\geq$ 80-nt residual tract with net excess $\geq 12$ (10 partial; compared length < 4,000 nt) and Fisher $p < 10^{-4}$.

Non-certified mosaics default to `URF`. Separately, any Tier-1 segment ($\geq$ 80 valid nt) falling below the pack's `min_segment_identity` (82% HIV-1, 90% IAV/SARS-CoV-2, 70% sarbecoviruses) or violating an active temporal floor is relabeled `OutOfPanel`, causing immediate `OutOfPanel` routing (whereas 100%-`Unattributed` single-segment queries return `Unclassifiable`). For HIV-1, `URF` outputs include a 25-trunk epidemiological trace (13 pure basal trunks excluding `?E` + 12 pandemic founder CRFs at complexity offset 0.005/site, plus `Unattributed`) rendered under the formal BNF grammar (Table S2). In SARS-CoV-2 surveillance streams, the post-2021 Wuhan-`B` amplicon-backfill QA check is applied as a lightweight post-processing audit filter over `EGGMAN` Tier-1 segment outputs in the Python streaming scripts.

##### S1.5 Benchmarking Protocols, External Comparators, and Phylogenetic Verification

External comparators ran on identical unaligned FASTA inputs: `COMET v2.4` [8] and 28-worker `jpHMM` [9] on 1,093 HIV-1 queries and 500 GTR + $\Gamma_4$ synthetic mosaics (900 true junctions); 4-thread `USDA GenoFLU` (`blastn` ≥ 98% allele matching) [2] on 1,200 SRA H5N1 and 650 masked `B3.13/D1.1` OOD isolates; and 8-thread `Nextclade CLI` [3] on 1,000 curated clinical SARS-CoV-2 genomes and 8 masked `XDV/XFZ/XFC` OOD isolates. Non-recombinant false-positive simulations (1,300 HIV-1 genomes across all 13 basal trunks) spanned baseline evolution, 3× rate acceleration, and APOBEC3G/3F G→A hypermutation. Orthogonal ML phylogenies for all surveillance findings and clock comparisons were inferred in `IQ-TREE 2` (GTR + $\Gamma_4$, 1,000 `UFBoot2` replicates) [10, 11] and dated with `LSD2` [12] and `TreeTime` [13].

##### S1.6 Census-Level Supply, Offspring Scaling, and Two-Stage Selective Sieve Equations

Across $R = 32$ countries ($n_r \geq 40$ deduplicated hosts), non-URF heterozygosity $H_{\text{circ},r} = 1 - \sum_m p_{r,m}^2$ was regressed against patient-level URF prevalence via Spearman rank correlation, sample-size WLS, heteroskedasticity-consistent binomial GLM (`HC3`), and offset negative-binomial GLM controlling for Sub-Saharan Africa, sample size, and sampling span. Offspring cluster sizes ($N = 761$ mosaic lineages) were fit by discrete MLE to Hurwitz-zeta ($P(N = k) = k^{-\alpha}/\zeta(\alpha, x_{\min})$), discrete lognormal (under both continuity-corrected $[k-0.5, k+0.5]$ and $[k, k+1)$ discretizations), Yule–Simon, and negative-binomial models, paired with Clauset parametric bootstrap goodness-of-fit (400 replicates) and Vuong likelihood-ratio tests [14], and checked for invariance across host deduplication thresholds (0.4%, 0.8%, and 1.5% Hamming divergence across all 31,825 full-length GenBank genomes). For evolutionary sieve two-locus tests across cohort $\mathcal{T}$ with genome parent shares $f_{g,k}$, expected binomial concordance and normalized $Z$-statistics follow:

$$\mathbb{E}_{\text{null}}(g) = \sum_k f_{g,k}^2, \qquad Z = \frac{\sum_{g \in \mathcal{T}} \left[Y_g - \mathbb{E}_{\text{null}}(g)\right]}{\sqrt{\sum_{g \in \mathcal{T}} \mathbb{E}_{\text{null}}(g)\left[1 - \mathbb{E}_{\text{null}}(g)\right]}}, \tag{6}$$

paired with 10,000 linear segment permutations, circular genome shifts, parental-trunk–stratified CMH odds ratios, multivariable logistic regression (adjusting for total breakpoint count and parental trunk pair), two `EGGMAN` Stage-1 detectability nulls (1,000 single-crossover mosaics at 2% noise plus 1,000 realistic 4-crossover mosaics at 6%–10% GTR + $\Gamma_4$ divergence), and explicit Stage-2 phylogenetic pseudoreplication controls restricting established CRFs to the 71 alias-collapsed Generation-1 primary founder CRFs alongside 527 DAG-deduplicated *de novo* CRF breakpoints.

#### Supplementary Text

##### Section S1: Basal Reference Scrubbing and the Locus-Restricted `?E` Trunk

Reference-free `RhizAeon` scanning [5] of the 2023 LANL pure-subtype references exposed 12 foreign or divergent mosaic intervals across 10 canonical genomes in HXB2 5885–9702 ($p \leq 1.83 \times 10^{-6}$, 7 involving unindexed donors; Table S3). Applying the explicit non-loop scrubbing rule (masking/demoting reproducible inter-subtype tracts spanning conserved non-loop genes while retaining isolated hypervariable *gp120 V1–V5* loop signals already masked in Viterbi), 7/55 (12.7%) pure references resolved directly as basal mosaics in `EGGMAN` Tier 1: all four Subtype G references (`DRCBL AF084936`, `HH8793 AF061641`, `92NG083 U88826`, and `PT2695 AY612637` carrying `G0[A1:gp41..nef]` [15]) and all three Subtype A7 references (`KX389622`, `KX389608`, `MH078558`), while `A3/A4/A8` collapsed into `A1` and Subtype L `AF286236` masked a 3′ `K:nef` tract (8976–9417) prior to consensus construction. Because `?E` contributed zero mosaic tracts upstream of HXB2 5097 across both LANL references and archive screens, locking `?E` to HXB2 5097–9417 ($\delta_{?\text{E}} = 0.035$) eliminated 5′ false switches [16, 17].

#### Section S2: Bridged Breakpoint Audit, Chronology, and Recombination DAG

Across 986 testable LANL CRF junctions obtained after bridging 106 ambiguous X/Y plateaus (80/106 = 75.5% verified statistically at compendium $\gamma = 10.5$), 493 (50.0%) represent verified elementary switches, 219 (22.2%) represent unexpanded second-generation CRF labels, 58 (5.9%) involve unclassified U donors untestable against an incomplete Tier-1 basis, and 216 (21.9% of 986, or 23.3% of 928 indexed-donor junctions) fail statistical support (90 unbridgeable X/Y [9.1%], 61 < 3-SNP floor [6.2%], 57 failed Fisher $p > 10^{-3}$ [5.8%], and 8 boundary artifacts [0.8%]). Nine shared mosaic structures explain 20 distinct CRFs (11 aliases), while three CRFs failed mosaic verification and were demoted alongside four Subtype A sub-subtypes (CRF26_A5U, CRF92_C2U, and CRF46_BF1; CRF31_BC, CRF35_A1D, and CRF146_BC remained active). The resulting 145-CRF genealogy DAG places 3 in Generation 0, 72 in Generation 1, 55 in Generation 2, and 15 in Generation 3, with 60/70 (85.7%) Generation 2–3 CRFs tracing transitive ancestry back to CRF01_AE (27), CRF07_BC (19), and CRF02_AG (17). Across 21 Group M clades (3,594 genomes; using a two-regime spline for Subtype B), full-clade ChronAeon divergence times correlated with LSD2 ($\rho = 0.814$, $p = 7.04 \times 10^{-6}$, median absolute difference 12.72 yr) and TreeTime ($\rho = 0.791$, $p = 1.97 \times 10^{-5}$, median difference 11.23 yr; $\rho = 0.808$ across all 27 clade/segment runs), with point estimates placing Subtype C (1951.4) and CRF02_AG (1956.1, 1938.2–1971.0) prior to Subtype A1 (1959.7) and Subtype G (1961.5, 1947.0–1973.9) within overlapping confidence intervals and clock uncertainty, followed by CRF01_AE (1974.7), A6 (1982.8), CRF07_BC (1988.2, open upper CI), CRF08_BC (1992.7), and CRF63_02A6 (2009.6).

#### Section S3: HIV-1 Benchmarking and the Stanford HIVDB Audit

Tables S4, S5, and S6 detail HIV-1 performance across 1,093 unaligned queries (0.510 s local, 2,144.1 seq/s; 2.919 s Apple M3 vs. 11,280.0 s wall / 271,492.9 s CPU for 28-worker jpHMM), 500 simulated GTR + $\Gamma_4$ mosaics (9.0 nt median error, 97.11% ≤ 100 nt, 7.87% false junctions vs. jpHMM 10.0 nt, 89.33%, 41.78% false junctions in 7,279.8 s wall), 536 combined LOSO runs (100.0% parent recall; 69.70% exact and 94.14% URF upon lineage omission across 495 CRFs), matched CRF01-118 OOD holdouts ($n = 152$: 141 URF + 3 OutOfPanel = 144/152 [94.74%] non-legacy abstention + 8/152 [5.26%] false legacy calls, tying COMET's 144/152 [94.74%] unassigned + 8/152 [5.26%] false legacy, with symmetric 25-trunk founder-CRF hints on 101/152 [66.4%] vs. COMET's 115/152 [75.66%] legacy-hinted calls, while exact independent LANL parent-set recovery on the 94 held-out CRF119-157 genomes reached 64/94 [68.1%, Jaccard 0.903] for EGGMAN Tier 1 vs. 13/94 [13.8%, Jaccard 0.628] for COMET), and 1,300 null simulations (0.0% Tier-1 FPR; 99.85% Tier-2 pure across standard, rate-spike, and APOBEC regimes). Evaluating all three tools symmetrically against independent published LANL CRF parental dictionaries (A-collapsed; excluding circular hotspot panels) confirmed out-of-sample clinical CRF parent-set superiority (94.6% [80.5% Tier-1 only], Jaccard 0.976 vs. 77.9% / 0.891 COMET and 77.9% / 0.919 jpHMM).

On Stanford HIVDB genotype-references.fasta ($N = 720$), EGGMAN matched 660/720 (91.7%: 473 pure + 187 CRF; 98.3% pure refs) and flagged six nominal pure refs with mosaic *pol* tracts (KC018549, KT022392, KT022405, KU820851, KX228808, KY658687). Across hiv-survdb ($N = 120{,}468$), partitioning 111,905 isolates.d-linked sequences from 8,563 unlinked FASTAs (newly subtyped: 6,410 pure, 1,569 CRF, 367 URF, 217 Subtype G) showed 88.38% concordance (72,680 pure + 26,221 CRF) while accounting for all 13,004 non-concordant linked records: 6,141 (5.49%) multi-enzyme locus splits (B|CRF07_BC $n = 2{,}396$), 2,157 (1.93%) true intra-patient clonal contradictions (CRF06_cpx|G $n = 520$ in Estonia), 1,453 CRFs with pure *pol*, 1,444 pure samples with mosaic *pol*, 1,243 resolved ambiguous tags (997 B + C), 486 alternate CRF/trunk/01_AE calls, and 80 (0.07%) short or unclassifiable fragments. Tier-2 clade guards rescued 190/218 (87.2%) basal G0[A6:RT] traces directly to CRF02_AG (40/40 West African ML trees closer to CRF02_AG: 0.0691 vs. 0.1560 subst/site to A6/CRF63).

#### Section S4: Three-Class GenBank Archive Audit of Full and Partial Mosaics

Deduplicating 31,825 full-length HIV-1 GenBank genomes (< 0.8% within identical barcodes) retained 7,720 independent patients alongside 20,797 strictly partial-only patients (7,897 *pol* across 10,365 sequences; 12,900 *env* across 27,885 sequences). Multi-segment `IQ-TREE 2` auditing separated unindexed clusters into (i) 8 blindly rediscovered post-2023 LANL CRFs (`CRF127`, `135`, `136`, `162`, `172`, `173`, `188`, `205`); (ii) 1 strictly monophyletic full-genome regional candidate, Russian `RU_G0B_gagMA` ($N = 6$, 2014–2016, 4.49% divergence, 4/4 segments at 100% UFBoot; Table S7 and Fig. S1A), plus 1/12 monophyletic partial-*env* cluster, Tanzanian `C[D:gp41]` ($N = 3$ patients across 3 independent studies, 8.26% divergence, 2/2 segments monophyletic and parent-concordant at 93%/98% UFBoot; unpromotable to CRF solely because full-genome coverage is absent per Robertson et al. [18]); and (iii) convergent polyphyletic hotspot URFs (or divergent single-subtype sub-lineages, as in `G0[H:gp41..nef]`, which is cluster-monophyletic on 3/3 segments [75% min UFBoot] but parent-trunk concordant on only 1/3) spanning 7 full-genome formulas (led by East African `D[A1:env]`, $N = 11$, *env* UFBoot 50%; Fig. S1B) plus 11/12 partial-gene hotspots (5 *pol*, 20 patients; 6 *env*, 81 patients; 101/104 total partial patients, led by `A1[D:gp41]` [$N = 49$, 15 studies, UFBoot 58%] and `A1[C:gp41]` [$N = 15$, 11 studies, 65%]; Table S8).

#### Section S5: Regional Heterozygosity, Deduplication Sensitivity, and Founder-Deduplicated Sieve Controls

Across $R = 32$ countries ($n_r \geq 40$), non-URF heterozygosity $H_{\text{circ}}$ predicted URF prevalence across Spearman ($\rho = 0.589, p = 3.86 \times 10^{-4}$), WLS ($R^2 = 0.621, p = 1.59 \times 10^{-4}$), `HC3` binomial GLM ($p = 1.45 \times 10^{-7}$), and negative-binomial GLM ($p = 3.54 \times 10^{-6}$). Offspring cluster sizes ($N = 761$) showed a steep heavy-tailed decline spanning three orders of magnitude ($N = 1$: 71.6%; $N \geq 3$: 19.1%; $N \geq 10$: 4.1%), approximated by a discrete Hurwitz-zeta tail ($\hat{\alpha} = 2.2095 \pm 0.0477$, AIC = 2015.37, KS = 0.0426; Yule–Simon $\Delta$AIC = 33.81), though Clauset bootstrap goodness-of-fit rejected a strict single-exponent power law from $x_{\min} = 1$ ($p < 0.003$, 0/400 replicates) and unconstrained discrete lognormal fit comparably or better depending on discretization ($[k-0.5, k+0.5]$: LL = −1002.60, AIC = 2009.19, Vuong $Z = -3.17, p = 0.0015$; $[k, k+1)$: AIC = 2049.64, Vuong $Z = +3.55$). Varying the host deduplication threshold across all 31,825 full-length GenBank genomes (0.4%, 0.8%, and 1.5% Hamming divergence) preserved both cluster-tail proportions ($N = 1$: 64.9%, 65.1%, 65.5%; $N \geq 3$: 21.9%, 21.4%, 21.2%; $N \geq 10$: 3.6%, 3.4%, 3.2%; $\hat{\alpha} = 2.13$–$2.15$) and Stage-2 singleton URF rates (`Intact 3' Block`: 22.1%, 22.0%, 22.2%, remaining significantly depleted vs. all established CRFs $p \leq 0.0051$ and Gen-1 primary CRFs $p \leq 0.029$; *RRE–rev2*: 64.1%–64.2%; *tat1–tat2*: 56.7%–56.8%).

Stage 1 breakpoint avoidance held across uniform, diagnostic-SNP, 2%-noise 1-BP detectability (1,204 full-screen / 1,222 default recalled BPs), and realistic 4-crossover 6%–10% (8% mean) GTR+$\Gamma_4$ detectability nulls (4,126/4,000 full-screen / 4,136/4,000 default recalled BPs) for both *gp120* core ($0.25\times$–$0.30\times$, $p \leq 1.58 \times 10^{-16}$; 0.27× under 4-BP 8%-div null) and *Gag*-CA core ($0.36\times$–$0.55\times$, $p \leq 0.0095$; combined two rigid cores 45/701 = 6.4% Est, $0.29\times$–$0.35\times$, $p \leq 3.3 \times 10^{-19}$; canonical *gag.CA* 1186–1878 0.46×, $p = 5.38 \times 10^{-6}$; *env.gp120* 6225–7757 0.63×, $p = 1.50 \times 10^{-6}$; Table S9), while the four inter-domain hinges absorbed 55.8%–60.8% of breaks (1.15× vs. 2% null, 1.47× vs. 4-BP 8% null). By contrast, *Pol*-RT (2650–3450) proved crossover-tolerant (11.7% Est; 63/527 = 12.0% DAG-deduplicated *de novo* CRF breakpoints vs. 6.7% URF, OR = 1.89, $p = 1.18 \times 10^{-4}$; 9.9% Gen-1, CMH $p = 0.15$). At Stage 2, eliminating Generation 2–3 CRF hub pseudoreplication by restricting established CRFs to the 71 alias-collapsed Generation-1 primary founder CRFs confirmed that point *tat1–tat2* (5950 ↔ 8400) remained non-significant everywhere (all-CRF CMH $p = 0.42$, MV $p = 0.16$; Gen-1 63.4%, CMH $p = 0.16$, MV $p = 0.093$), while preserving the undivided 3′ regulatory/Env cassette (HXB2 5831–8653: 17.9% [88/491] URF → 33.1% [50/151] all Est, CMH OR = 2.20, $p = 1.15 \times 10^{-3}$, MV [adj. $n_{\text{bp}}$ + pair] OR = 3.09

[95% CI 1.87–5.09], $p = 9.61 \times 10^{-6}$; Gen-1 dedup 33.8% [24/71], CMH OR = 2.11, $p = 0.025$, MV OR = 2.58 [1.35–4.92], $p = 3.96 \times 10^{-3}$). Within that 3′ block, *RRE–rev2* strengthened in Gen-1 founders (81.7% [58/71], exceeding its own 73.6% Gen-1 linear null, $p = 0.046$; CMH OR = 2.31, $p = 0.021$; MV OR = 2.67 [1.37–5.20], $p = 4.01 \times 10^{-3}$; all-CRF 72.2%, MV $p = 0.027$ adjusting for $n_{bp}$ + pair, attenuating to $p = 0.119$ when conditioning on colinear 8150–8550 breaks). Conversely, crude suppression of *gp41*-CT return swaps among foreign-*gp120* lineages (110/309 = 35.6% URF vs. 17/87 = 19.5% all Est, crude $p = 0.0042$) vanished under parental-pair CMH (OR = 0.79, $p = 0.60$) and Gen-1 deduplication (8/34 = 23.5%, CMH OR = 0.86, $p = 0.97$), and upstream *gag/pol* enrichment (7.0% → 10.7% → 15.1% major; 51/527 = 9.7% DAG-dedup *de novo* BPs, $p = 0.042$) attenuated in Gen-1 alone (9.3%, $p = 0.16$).

#### Section S6: Multi-Pathogen Streaming, Head-to-Head Comparators, and Temporal Guards

Table S10 summarizes all cross-family benchmarks. In Influenza A H5N1 (1,200 SRA isolates, 0.127 s), `EGGMAN` scored 97.82% (1,076/1,100) across all 10 USDA-labeled genotypes (149/150 pure + 927/950 reassortants including 95/100 `C2.1`, with zero cross-genotype misclassifications; 23 `OutOfPanel` + 1 `URF` safe abstentions; 0.127 s vs. 3.230 s `GenoFLU`), assigned structured calls to 98% of 100 USDA-unassigned assemblies (52 certified, 25 `URF`, 21 `OutOfPanel`; 92% multi-segment mosaics; 99% mosaic rate across the companion unassigned subset with 16/16 ML trees at 69%–100% UFBoot), and flagged 100% of 650 masked `B3.13/D1.1` OOD isolates (98.92% exact donor set in 0.089 s vs. 4-thread `GenoFLU` 100% `Not_assigned` in 1.712 s, a 19.2× wall / 77× CPU speedup). On 3,980 BV-BRC H5Nx assemblies (7.311 s, 544.4 iso/s; 3,966 ≥ 11.5 kb), 420 `URF`s separated into 20 constellations (352 isolates: 12 omitted USDA/GenoFLU across 114 isolates; 8 EURL/non-USDA incl. `euDI.2.1` across 238 isolates) plus 68 rare isolates. On curated SARS-CoV-2 ($N = 1{,}000$, frozen 16-founder pack), default thresholds ($\gamma = 7.0$, $m_{\min} = 5$, $p < 10^{-4}$; 0.592 s) achieved 100.0% Tier-1 pure specificity, 78.5% Tier-1 state accuracy, 96.4% combined state accuracy, and 80.7% Tier-2 exact accuracy; CoV-tuned thresholds ($\gamma = 3.5$, $m_{\min} = 3$, $p < 0.01$; 1.059 s including wrapper overhead vs. 3.424 s 8-thread `Nextclade`) achieved 98.2% (491/500) Tier-1 pure specificity (99.4% exact pure waves), 81.0% (405/500) Tier-1 recombinant breakpoint detection (89.6% combined Tier-1 state), 97.3% combined clonal/recombinant accuracy (vs. `Nextclade` 98.3%), and 80.4% Tier-2 exact family concordance (146/157 `XBB` `URF`s keeping exact Tier-1 coordinates). On independent BV-BRC streams, the 16-founder basis cut pure-wave flags eightfold (5.96% → 0.72% CoV-tuned; 0.09% default across 13,275 pure-wave genomes in 23,376 total), scanned 100,000 live genomes in 113.1 s default / 118.3 s CoV-tuned, flagged 3/79 Wuhan-`B` backfill QA hits via post-processing audit filters, and recovered 13 CoV-tuned `URF` clusters (1,434 genomes: 10/13 = 1,401 [97.7%] matching 10 Pango `X*` roots). All 79 valid sarbecoviruses ran in 0.066 s (5/5 clonal basals, 62/74 = 83.8% mosaic). Across all 16 temporal-guard runs (140,276 genomes), `--temporal-guard` (`--temporal-mode sampled|submission`) achieved zero collection- and submission-date floor violations.

### Supplementary Figures

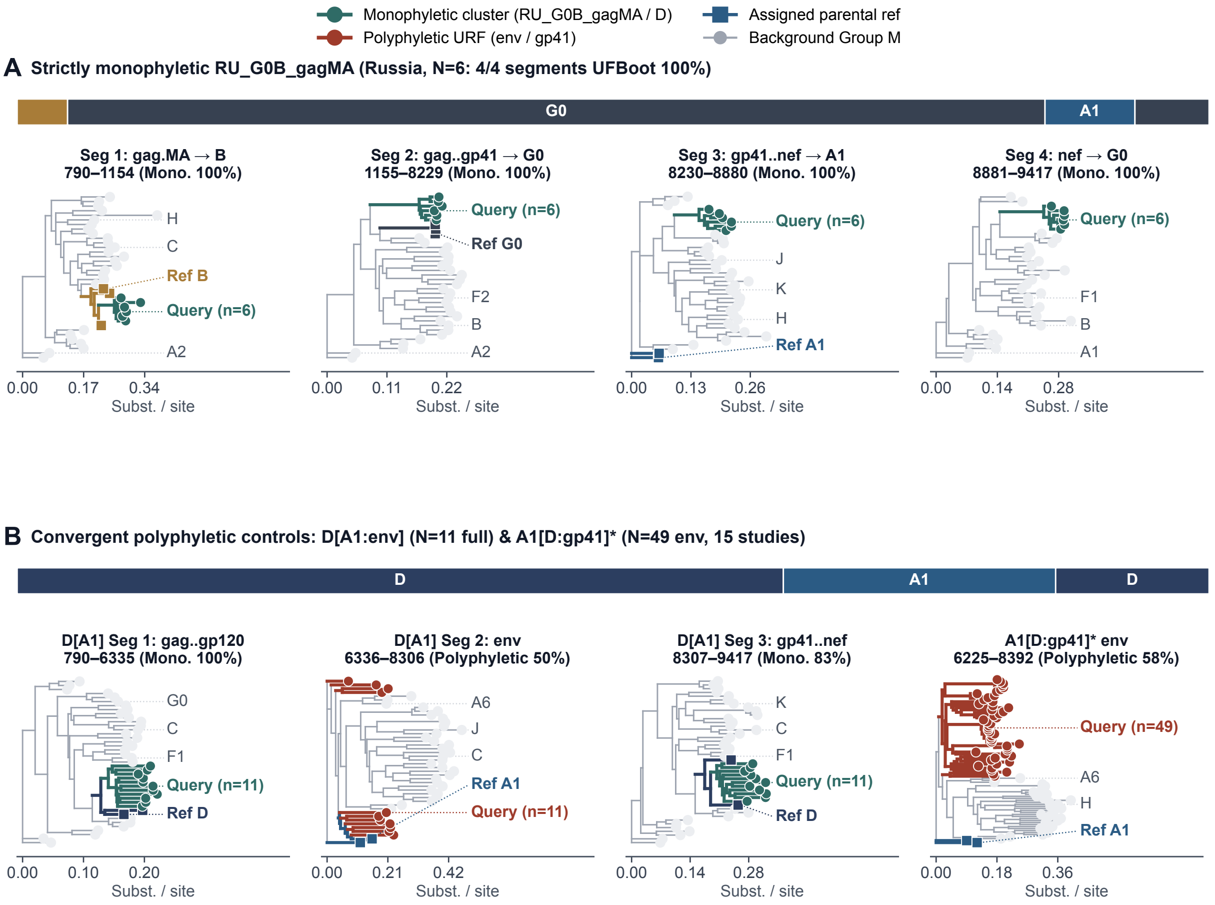


Figure S1: **Maximum-likelihood segment phylogenies separating monophyletic candidate CRF `RU_G0B_gagMA` from convergent polyphyletic HIV-1 hotspots.** Midpoint/rooted IQ-TREE 2 phylograms (GTR + $\Gamma_4$, 1,000 `UFBoot2` replicates; scale bars, substitutions per site) highlight monophyletic query clusters in teal, polyphyletic hotspot queries in vermilion, assigned parental references as blue squares, and Group M background in gray, with true-coordinate HXB2 genome diagrams above each row. **A:** Russian candidate lineage `RU_G0B_gagMA` ($N = 6$, `[Span:2014-2016]`) forms a strictly monophyletic clade (100% UFBoot) across all four non-recombinant tracts: *gag.MA* → `B` (790–1154, 100%), *gag..gp41* → `G0` (1155–8229, 100%), *gp41..nef* → `A1` (8230–8880, 100%), and *nef* → `G0` (8881–9417, 100%). **B:** Convergent polyphyletic controls combine full-genome East African `D[A1:env]` ($N = 11$, monophyletic on backbone *gag..gp120* `D` [100%] and 3′ *gp41..nef* `D` [83%] but polyphyletic across multiple independent `A1` captures inside *env* 6336–8306 [50% UFBoot]) with partial-*env* hotspot `A1[D:gp41]*` ($N = 49$ patients across 15 studies, polyphyletic across `A1` 6225–8392 [58% UFBoot]).

### Supplementary Tables

Table S1: First-principles parameterization of coordinate alignment, unit-mismatch Viterbi segmentation, and four-gate lineage calling in EGGMAN and ChronAeon.

| Parameter | Value | Derivation / Implementation Rule |
|---|---|---|
| Seed length $k$ | 15 nt (fallback 11 nt) | Random collision $4^{-15} = 9.31 \times 10^{-10} \ll 1/L$ (step 2 nt; 11-nt step 3 nt sparse rescue). |
| Bitmask NW | $+2/-1/-3$ linear; $B = 45$ nt | 4-bit A/C/G/T reference-union bitmask scoring within 45-nt band (query ambiguities skipped in Viterbi); split-DP for indels $> 180$ nt. |
| Viterbi emissions | 0.0/1.0; gap 0.35; Unattr 0.58 | Mismatch-equivalent (Hamming) units (2.0 out-of-range); Unattributed acts as a high-divergence / alignment-failure floor ($> 58\%$ mismatch). |
| Switch penalty $\gamma$ | 7.0 (10.5 / 13.5 / 3.5) | Divergence-calibrated mismatch-unit switch cost (1 mismatch $\approx \ln[(1-p)/(p/3)]$ nats: $\approx$ 3.5 nats HIV-1 vs. $\approx$ 8.0 nats SC2); segmented $\gamma = 0.0$ / intra $10^6$. |
| Unsampled ?E offset | $\delta_{?E} = 0.035$ /site, $x \geq 5097$ | Entropy equalization across *vif–nef*; zero emission support upstream of HXB2 5097. |
| Fisher boundary test | $p < 10^{-4}$ ($< 10^{-3}$ comp.) | Upper-tail 2×2 Fisher exact test across flanking spans ($\geq$ 120 nt full / 150 nt partial; $m_{\text{min}} \geq 5$ SNPs). |
| ChronAeon decay | $t_{1/2} = 6.93/0.50/0.25$ yr | Exponential weighting ($\lambda = 0.10$ yr$^{-1}$ HIV-1) with $\frac{1}{2}h_{QQ}$, $\alpha_\Gamma = 0.20$, and slack $\delta_t = 3.0/0.5$ yr (sampled) or 0.0 yr (submission). |
| Host deduplication | $d < 0.8\%$ within barcode | Collapses longitudinal / clonal amplification replicates prior to archive clustering. |
| Epi fallback offset | 0.005/site (25 named trunks) | Complexity penalty applied to 12 pandemic founder CRFs alongside 13 pure basal trunks (excl. ?E) + Unattributed. |
| Tier-2 four-gate floor | Id $\geq$ 84%; Stem $\Delta \geq 8/4/2$; Kadane $\geq$ 12 | Covers 152 active HIV-1 callers (171 catalogued): Gate 1 plurality, Gate 2 DAG dominance, Gate 3 stem $B_c(x)$ (8 HIV / 4 IAV / 2 SC2 full), Gate 4 Kadane $\geq$ 80 nt; OutOfPanel on any segment $<$ 82%/90%/70% id. |

Table S2: Formal Backus–Naur Form (BNF) grammar generating canonical descriptive coordinate barcodes across contiguous and segmented viral packs.

<table>
<tr><th>Nonterminal</th><th>Production Rule</th><th>Semantic Meaning</th></tr>
<tr><td><barcode></td><td><backbone> "[" <donor_list> "]" <qual> | <backbone>"(Clonal)" <qual></td><td>Recombinant donor structure or non-mosaic trunk with optional scope/catalog qualifier.</td></tr>
<tr><td><backbone></td><td><trunk> | <founder_crf></td><td>Plurality basal parental trunk or epidemiologically certified founder backbone.</td></tr>
<tr><td><donor_list></td><td><donor_tract> | <donor_tract> "|" <donor_list></td><td>Pipe-delimited set of minority donor tracts across the genome.</td></tr>
<tr><td><donor_tract></td><td><trunk> ":" <span_list></td><td>Single parental donor trunk paired with its coordinate-indexed gene spans.</td></tr>
<tr><td><span_list></td><td><span_item> | <span_item> "+" <span_list></td><td>Disjoint homologous or reassorted tracts derived from the same parental donor trunk.</td></tr>
<tr><td><span_item></td><td><locus> | <locus> ".." <locus></td><td>Single genomic locus/segment or contiguous gene/domain range.</td></tr>
<tr><td><locus></td><td>"gag.MA" | "PR" | "PB2" | "HA" | "S" | ...</td><td>Standardized genomic landmark or RNA segment anchored to pack reference coordinates.</td></tr>
<tr><td><qual></td><td>[<scope>] [<epoch>]</td><td>Optional dynamic amplicon scope and catalog-annotated temporal qualifier.</td></tr>
<tr><td><scope></td><td>"@pol" | "@env" | ""</td><td>Partial-genome amplicon projection (emitted dynamically; omitted for full genomes).</td></tr>
<tr><td><epoch></td><td>"@" <year> | "[Span:" <year> ["-" <year>] "]"</td><td>Catalog-level ChronAeon $t_{\text{MRCA}}$ or observed sampling span annotation.</td></tr>
</table>

Table S3: Twelve verified divergent or inter-subtype mosaic tracts identified across ten nominal pure-subtype LANL references in reference-free `RhizAeon` scanning [5] (HXB2 5885–9702).

| Nominal Label | Accession | Declared Subtype | Verified Tract (HXB2 Span) | Fisher $p$ |
|---|---|---|---|---|
| A8_CV_10_115 | MW353966 | Subtype A8 | *vpu..3'LTR* (6056–9702; Unindexed in A1) | $1.87 \times 10^{-15}$ |
| C_IN_95_95IN21068 | AF067155 | Subtype C | *env-gp120(V1–V3)* (6602–6741; Unindexed in C) | $4.52 \times 10^{-14}$ |
| C_IN_95_95IN21068 | AF067155 | Subtype C | *env-gp120(C3–V5)* (7233–7438; A2 in C) | $7.94 \times 10^{-8}$ |
| B_TH_90_BK132 | AY173951[b] | Subtype B | *env-gp120(V1–V5)* (6599–7764; Unindexed in D) | $1.74 \times 10^{-13}$ |
| B_TH_90_BK132 | AY173951[b] | Subtype B | *env-gp120–3'LTR* (7146–9702; Unindexed in F2) | $6.93 \times 10^{-8}$ |
| † A7_NG_11_DEURF | MH078558 | Subtype A7 | *env-gp120–gp41* (6316–8301; Unindexed in A7) | $2.14 \times 10^{-13}$ |
| A2_CY_94_94CY017 | AF286237 | Subtype A2 | *env-gp120–3'LTR* (6621–9702; Unindexed in A2) | $2.43 \times 10^{-13}$ |
| H_CF_90_056 | AF005496 | Subtype H | *env-gp120(C3–V5)* (7283–7463; Unindexed in H) | $1.36 \times 10^{-10}$ |
| G_PT_x_PT2695 | AY612637 | Subtype G[a] | *env-gp120–nef* (6627–8880; A1 in G) | $2.42 \times 10^{-10}$ |
| F1_BR_93_93BR020 | AF005494 | Subtype F1 | *env-gp120(V1–V5)* (7074–7651; G in F2) | $3.43 \times 10^{-10}$ |
| L_CD_83_83CD003 | AF286236 | Subtype L | *nef* (8976–9417; K in L) | $1.74 \times 10^{-6}$ |
| A3_SN_01_DDJ369 | AY521631 | Subtype A3 | *vpr–tat* (5885–6054; G in A7) | $1.83 \times 10^{-6}$ |

[a] When evaluated against scrubbed G0 and A1 profiles, all four nominal Subtype G references (AF084936, AF061641, AY612637, U88826) exhibit the shared G0[A1:gp41..nef] ancestral mosaic. [b] Asian Subtype B AY173951 resolves cleanly into clonal B under multi-trunk scrubbed emissions. † EGGMAN Tier 1 across $K = 14$ profiles decomposes 7/55 (12.7%) nominal pure references directly into non-clonal mosaics (all four Subtype G + three demoted Subtype A7), while A3/A4/A8 map to A1 and L masks K:nef.

Table S4: Head-to-head benchmark of `EGGMAN` versus `COMET v2.4` across 1,093 unaligned HIV-1 sequences spanning twelve evaluation panels and strata.

| Benchmark Panel | $N$ | CM Exact | CM Unass. | Pure | CRF | URF | Comp. | EG Primary | Basal BP |
|---|---|---|---|---|---|---|---|---|---|
| 1A. Pure Subtype Refs | 55 | 55 (100.0%) | 0 (0.0%) | 52 | 0 | 3 | 45 (81.8%) | 81.8% Exact | 7 (12.7%) |
| 1B. CRF01-CRF118 Refs | 369 | 362 (98.1%) | 6 (1.6%) | 8 | 347 | 14 | 359 (97.3%) | 94.0% Exact | 360 (97.6%) |
| 1C. CRF119-CRF157 Refs | 94 | 0 (0.0%) | 92 (97.9%) | 0 | 88 | 6 | 92 (97.9%) | 93.6% Exact | 93 (98.9%) |
| 1BC. All 145 LANL CRFs | 463 | 362 (78.2%) | 98 (21.2%) | 8 | 435 | 20 | 451 (97.4%) | 94.0% Exact | 453 (97.8%) |
| 1BC. Active 142 CRFs | 454 | 353 (77.8%) | 98 (21.6%) | 0 | 435 | 19 | 451 (99.3%) | 95.8% Exact | 452 (99.6%) |
| 2. Audited Panel 2 Total | 61 | 23 (37.7%) | 52 (85.2%) | 0 | 34+6 nov. | 21 | 54 (88.5%) | 86.9% Exact | 61 (100.0%) |
| 2a. Post-2023 + CRF63 | 31 | 3 (9.7%) | 26 (83.9%) | 0 | 29 | 2 | 30 (96.8%) | 93.5% Exact | 31 (100.0%) |
| 2b. RU_G0B_gagMA | 6 | 0 (0.0%) | 6 (100.0%) | 0 | 6 nov. | 0 | 6 (100.0%) | 100.0% Novel | 6 (100.0%) |
| 2c. Polyphyletic URFs | 24 | 20 (83.3%) | 20 (83.3%) | 0 | 5 | 19 | 18 (75.0%) | 79.2% URF | 24 (100.0%) |
| 3A. Partial *pol* Hotspots | 53 | 0 (0.0%) | 32 (60.4%) | 0 | 17 | 36 | 53 (100.0%) | 67.9% URF | 53 (100.0%) |
| 3B. Partial *env* Hotspots | 61 | 0 (0.0%) | 59 (96.7%) | 1 | 0 | 60 | 61 (100.0%) | 98.4% URF | 60 (98.4%) |
| 4A. Clinical Pure Subtypes | 150 | 141 (94.0%) | 9 (6.0%) | 147 | 0 | 3 | 147 (98.0%) | 98.0% Pure | 13 (8.7%) |
| 4B. Clinical Named CRFs | 150 | 107 (71.3%) | 43 (28.7%) | 1 | 108 | 41 | 139 (92.7%) | 72.0% Exact | 147 (98.0%) |
| 4C. Clinical URFs | 100 | 0 (0.0%) | 99 (99.0%) | 6 | 3 | 91 | 90 (90.0%) | 91.0% URF | 94 (94.0%) |

Exact COMET match on 2c counts unassigned calls on polyphyletic URFs matching expected URF ground truth; in strict named-CRF accuracy (Table S5), COMET scores 0.0% on Panel 2. Wrong CRF calls by EGGMAN were 0 across 1A, 1C, 1BC-active, 2a, 2b, and 4A, and 1/150 in 4B.

Table S5: Three-way comparison of `EGGMAN`, `COMET v2.4`, and `jpHMM` across 1,093 unaligned HIV-1 sequences (all 1,093 parsed) and 500 synthetic GTR + $\Gamma_4$ mosaics, with symmetric non-circular parental-set evaluation against independent published LANL CRF dictionaries.

| | | Exact Lineage (%) | | | Mosaic / Pure State (%) | | | Exact Parent Set (%)‡ | | |
|---|---|---|---|---|---|---|---|---|---|---|
| Benchmark Panel | *N* | EG | CM | JP† | EG | CM | JP | EG (T1) | CM | JP |
| 1A. Pure Subtype Refs | 55 | 81.8 | 100.0 | 85.5 | 81.8 | 100.0 | 85.5 | 87.3 (87.3) | 100.0 | 85.5 |
| 1B. `CRF01-CRF118` Refs§ | 369 | 94.0 | 98.1 | — | 95.9 | 98.1 | 98.4 | 97.0 (75.1) | 98.9 | 80.5 |
| 1C. `CRF119-CRF157` Refs | 94 | 93.6 | 0.0 | — | 98.9 | 97.9 | 100.0 | 96.8 (68.1) | 13.8 | 79.8 |
| 1BC. All 145 LANL CRFs§ | 463 | 94.0 | 78.2 | — | 96.5 | 78.2 | 98.7 | 96.9 (73.6) | 81.5 | 80.4 |
| 1BC. Active 142 CRFs§ | 454 | 95.8 | 77.8 | — | 98.2 | 77.8 | 99.6 | 97.8 (74.0) | 81.1 | 80.4 |
| 2. Audited Panel 2 | 61 | 86.9 | 0.0 | — | 95.1 | 85.2 | 100.0 | — | — | — |
| 3A. Partial *pol* Hotspots | 53 | 0.0 | 0.0 | — | 67.9 | 60.4 | 100.0 | — | — | — |
| 3B. Partial *env* Hotspots | 61 | 0.0 | 0.0 | — | 98.4 | 96.7 | 88.5 | — | — | — |
| 4A. Clinical Pure Subtypes | 150 | 98.0 | 94.0 | 92.0 | 98.0 | 94.0 | 92.0 | 92.0 (92.0) | 94.7 | 92.0 |
| 4B. Clinical Named CRFs§ | 150 | 72.0 | 71.3 | — | 97.3 | 71.3 | 99.3 | 94.6 (80.5) | 77.9 | 77.9 |
| 4C. Clinical URFs | 100 | 0.0 | 0.0 | — | 90.0 | 99.0 | 97.0 | — | — | — |

Runtime (1,093 seqs): EG 0.510 s local (2,144.1 /s) / 2.919 s M3 (374.4 /s) vs. 28-worker JP 11,280.0 s wall (271,492.9 s CPU = 248.4 s/seq). Synthetic 500-mosaic GTR + $\Gamma_4$ benchmark (900 true junctions): EG (0.133 s, 3,764 /s) median error 9.0 nt (IQR 4.0–22.2), 97.11% ≤ 100 nt, 99.78% ≤ 250 nt, 7.87% false junctions, 91.4% exact parent set (Jaccard 0.975) vs. JP (7,279.8 s wall / 155,084.7 s CPU) 10.0 nt (IQR 3.0–32.0), 89.33% ≤ 100 nt, 94.67% ≤ 250 nt, 41.78% false junctions, 51.6% exact parent set (Jaccard 0.725). Across 483 unbridged LANL segments, EG median offset 292.0 nt vs. JP 89.0 nt. †`jpHMM` outputs subtype intervals rather than CRF labels (with standard A-collapsed pure-subtype scoring). ‡Independent parent-set accuracy maps every tool's output symmetrically via published LANL CRF parent dictionaries (A-collapsed; excluding circular GenBank panels 2, 3A, 3B and unannotated 4C); EG shows combined Tier-1+Tier-2 (Tier-1 coordinate-only in parentheses). Mean Jaccard (EG / CM / JP): 1A 0.920/1.000/0.884; 1B 0.989/0.995/0.930; 1C 0.990/0.628/0.927; 1BC-active 0.994/0.919/0.930; 4A 0.960/0.966/0.956; 4B 0.976/0.891/0.919. §Excludes 4 LANL sequences in 1B ($n$ = 365 / 450 active) and 1 in 4B ($n$ = 149) whose CRF lacks an independent LANL indexed parent set (e.g., `U`-only or unmapped legacy tag).

Table S6: Strict out-of-sample generalization and null false-positive benchmarks for `EGGMAN`: LOSO cross-validation ($N$ = 536), matched `CRF01-118` OOD holdouts ($N$ = 152), and simulated non-recombinant nulls ($N$ = 1,300 per parameter regime).

| Evaluation Regime | Stratum ($N$) | Exact / Pure | Non-Legacy Abst. | False Legacy / Hints | Indep. Parent Set |
|---|---|---|---|---|---|
| LOSO Basal LOO | 12 Basal Trunks (41) | 37 (90.24%) | 4 `URF` (9.76%) | 0 | 41/41 (100.0%)§ |
| LOSO Certified | Leave-One-Out (495) | 345 (69.70%) | 135 `URF` (27.27%) | 15 sibling aliases | 495/495 (100.0%)§ |
| Combined LOSO | Panels 1A–1C + 2 (536) | 382 (71.27%) | 139 `URF` (25.93%) | 15 sibling aliases | 536/536 (100.0%)§ |
| Lineage Omitted | Unseen Dropped (495) | — | 466 `URF` (94.14%) | 29 (5.86%) | 495/495 (100.0%)§ |
| Matched OOD (`01-118`) | `COMET v2.4` on 1C+2 (152) | 0.0% (OOD) | 144 unass. (94.74%)† | 8 (5.26%) + 107 hint† | 13/94 (13.8%); J 0.628‡ |
| Matched OOD (`01-118`) | `EGGMAN` on 1C+2 (152) | 0.0% (OOD) | 141 `URF`+3 OOP (94.74%) | 8 (5.26%); 101 trace† | 64/94 (68.1%); J 0.903‡ |
| Null Pure Sims | Default ($\gamma$ = 7.0, 1,300) | 1,298 (99.85%) | 2 `URF` (0.15%) | 0 (0.0% T1; 0.0% CRF) | 1,300/1,300 (100.0%) |
| Null Pure Sims | Partial/Full/Comp (1,300 × 3) | 1,298 (99.85%) | 2 `URF` (0.15%) | 0 (0.0% T1; 0.0% CRF) | 1,300/1,300 (100.0%) |

†Both tools tie at 144/152 (94.74%) total non-legacy abstention and 8/152 (5.26%) outright false legacy calls (`EGGMAN`: 141 `URF` + 3 `OutOfPanel` + 7 false CRF + 1 false pure; `COMET`: 144 `unassigned` + 8 false legacy). Symmetric optional ancestral hints: `COMET` attaches legacy-CRF hints to 107/144 unassigned calls (115/152 = 75.66% legacy-hinted overall; 37/152 = 24.34% hint-free), while `EGGMAN`'s 25-trunk trace notes an ancestral founder CRF on 101/152 (66.4%). ‡Exact independent LANL parental-set accuracy (and mean Jaccard) evaluated strictly on the 94 held-out `CRF119-157` sequences with published LANL parent dictionaries (`EGGMAN` Tier-1 OOD coordinates vs. `COMET`; ≥ 1-trunk overlap is 94/94 for both). §LOSO parent-recall checks ≥ 1 shared parental trunk against held-out profile. Null simulations span 650 standard GTR + $\Gamma_4$, 325 3× *pol/gp120* rate-spike (2 conservative `URF` rejects in Tier 2), and 325 APOBEC3G/3F bursts across all 13 non-`?E` trunks (0.53–0.54 s).

Table S7: Audited full-genome GenBank mosaic clusters ($N_{host} \geq 3$) separating monophyletic regional candidate RU_G0B_gagMA from seven polyphyletic convergent hotspot URFs (or divergent single-subtype sub-lineages), alongside eight blindly rediscovered post-2023 LANL CRFs.

| Status / Formula | Countries | Representative HXB2 Barcode | Span | $N_{host}$ | ML Segs | Min UFB |
|---|---|---|---|---|---|---|
| **Candidate Regional CRF** | | | | | | |
| G0[B:gag.MA \| A1:gp41..nef] | Russia (6) | B:790-1154 \| G0:1155-8229 \| A1:8230-8880 \| G0:8881-9417 | 2014–2016 | 6 | 4/4 mono | 100.0% |
| **Non-CRF / Convergent URFs** | | | | | | |
| D[A1:env] | UG (8), KE (2), TZ (1) | D:790-6335 \| A1:6336-8306 \| D:8307-9417 | 1993–2015 | 11† | 2/3 mono | 50.0% |
| G0[H:gp41..nef] | CU (1), KE (1), CM (1) | G0:790-8261 \| H:8262-8820 \| G0:8821-9417 | 1999–2011 | 3 | 3/3 (1/3 conc.)‡ | 75.0% |
| B[F1:IN..tat1+gp120..nef] | Uruguay (3) | B:790-4929 \| F1:4930-5941 \| B:5942-6374 \| F1:6375-9417 | 2012–2018 | 3 | 3/4 mono | 100.0% |
| C[A1:env] | KE, RW, ZA | C:790-6332 \| A1:6333-8250 \| C:8251-9417 | 2003–2019 | 3 | Distinct BPs (2 hom.) | |
| C[A1:nef] | TZ (2), SE (1) | C:796-8783 \| A1:8784-9412 | 2004–2008 | 3 | Distinct BPs (2 hom.) | |
| D[A1:gp120..gp41] | Uganda (3) | D:790-6614 \| A1:6615-8255 \| D:8256-9417 | 2009–2012 | 3 | Distinct BPs (2 hom.) | |
| F1[B:RT+tat1..gp41] | Argentina (3) | F1:796-2810 \| B:2811-3679 \| F1:3680-5941 \| B:5942-8623 \| F1:8624-9417 | Undated | 3 | Distinct BPs (1 hom.) | |

Note: Eight additional full-genome GenBank clusters first surfaced blindly mapped identically to newly official post-2023 LANL CRFs: CRF127_07109, CRF135_0107, CRF136_0107, CRF162_cpx, CRF172_0755, CRF173_63A6 (scrubbed from CRF63_02A6), CRF188_0107, and CRF205_0107. †Co-occurs with partial-*env* hotspot A1[D:gp41] ($N$ = 49 across 15 studies). ‡Cluster-monophyletic across 3/3 segments (75% min UFBoot), but parent-trunk concordant on only 1/3 (seg 2 assigned H nests sister to G0 at 85%; seg 3 assigned G0 nests sister to ?E at 71%), indicating a divergent Subtype G sub-lineage rather than a G0+H recombinant.

Table S8: All twelve partial-gene HIV-1 mosaic clusters ($N_{host} \geq 3$) identified across 20,797 strictly partial-only GenBank+LANL hosts (7,897 *pol* [10,365 seqs]; 12,900 *env* [27,885 seqs]): 1/12 (C[D:gp41]) is strictly monophyletic across both partial *env* segments (unpromotable solely because full-genome sequence is absent), whereas 11/12 (101/104 patients) classify phylogenetically as convergent polyphyletic hotspot URFs.

| Region | Window Formula | Representative HXB2 Barcode | Primary Countries | $N_{pat}$ | $N_{stud}$ | Mono Segs | Min UFB |
|---|---|---|---|---|---|---|---|
| *env* | A1[D:gp41] | A1:6225-8392 \| D:8393-8795 | UG (37), KE (9), TZ (3) | 49 | 15 | 0/2 | 58.0% |
| *env* | A1[C:gp41] | A1:6225-8397 \| C:8398-8771 | RW (6), MW (3), UG (2), TZ (1) | 15 | 11 | 0/2 | 65.0% |
| *env* | B[C:gp41] | B:6225-8084 \| C:8085-8795 | BR (2), ZA (2), IN (2) | 7 | 4 | 0/2 | 79.0% |
| *pol* | C[B:PR..RT] | C:2085-2374 \| B:2375-3181 \| C:3182-5096 | CN (7) | 7 | 3 | 1/3 | — |
| *env* | A1[C:gp120+gp41] | C:6225-6614 \| A1:6615-8160 \| C:8161-8795 | TZ (2), Unk/x (2) | 4 | 4 | 0/3 | 81.0% |
| *pol* | D[A1:IN] | D:2268-4268 \| A1:4269-5093 | UG (3), TZ (1) | 4 | 3 | 1/2 | 57.0% |
| *env* | C[D:gp41] | C:6225-7853 \| D:7854-8795 | TZ (3) | 3 | 3 | 2/2† | 93.0% |
| *env* | C[A2:gp41] | C:6225-8255 \| A2:8256-8795 | Unk (1), CD (1), ZM (1) | 3 | 2 | 0/2 | 80.0% |
| *env* | C[A1:gp120..gp41] | C:6225-7055 \| A1:7056-8250 \| C:8251-8795 | TZ (3) | 3 | 2 | 2/3 | 57.0% |
| *pol* | D[A1:gag..PR] | A1:2085-2517 \| D:2518-5096 | UG (3) | 3 | 2 | 1/2 | 61.0% |
| *pol* | C[D:gag..PR] | D:2085-2534 \| C:2535-5093 | ZA (3) | 3 | 2 | 1/2 | 100.0% |
| *pol* | A1[D:RT..IN] | A1:2292-4169 \| D:4170-5066 | UG (3) | 3 | 2 | 0/2 | 91.0% |

†Tanzanian C[D:gp41] ($N$ = 3 epidemiologically unlinked patients across 3 independent Tanzanian studies; 8.26% mean divergence) is strictly monophyletic and parent-trunk concordant across both partial *env* segments (C: 93%/100% UFBoot; D: 98%/99% UFBoot), remaining unpromotable to an official CRF solely due to the absence of full-genome sequence coverage per Robertson et al. [18] criteria. The remaining 11/12 partial clusters (101/104 patients) are polyphyletic convergent hotspots.

Table S9: Two-stage structural and epistatic sieve across 491 singleton URFs (2,046 BPs), 151 established CRFs (701 BPs), 22 major CRFs (93 BPs), and 71 alias-collapsed Generation-1 primary founder CRFs (plus 527 DAG-deduplicated *de novo* CRF breakpoints).

| Stage | Feature (HXB2 Coordinates) | Null† | URF | Est. / Maj. (G1) | Stage 1 / Stage 2 & Founder-Dedup Statistic |
|---|---|---|---|---|---|
| Stage 1 | *gp120* Core (6450–7650) | 13.9% (15.7/15.1) | 5.6% | 4.0% / 3.2% | 0.25× sim2 ($p = 7.4 \times 10^{-23}$); 0.27× sim4; 0.30× SNP |
| Stage 1 | *Gag*-CA Core (1280–1850) | 6.6% (4.4/6.7) | 3.1% | 2.4% / 3.2% | 0.55× sim2 ($p = 0.0095$); 0.36× sim4; 0.50× SNP |
| Stage 1 | Both Rigid Cores (*gp120+CA*) | 20.5% (20.1/21.8) | 8.7% | 6.4% / 6.5% | 0.32× sim2 ($p = 2.8 \times 10^{-24}$); 0.29× sim4 ($p = 7.1 \times 10^{-29}$) |
| Stage 1 | 4 Inter-Domain Hinges‡ | 34.8% (48.7/37.9) | 60.8% | 55.8% / 60.2% | 1.15× sim2 ($p = 1.8 \times 10^{-4}$); 1.47× sim4 ($p < 10^{-18}$) |
| Stage 1 | *gp120* ↔ *gp41* (6900 ↔ 7950) | 63.2% lin | 85.5% | 90.7% / 95.5% | $p_{\text{lin}} < 10^{-4}$; CMH OR = 2.00 ($p = 0.032$) |
| Stage 2 | Intact 3′ Block (5831–8653) | — | 17.9% | 33.1% / 36.4% (33.8%) | All: CMH OR = 2.20 ($p = 1.2 \times 10^{-3}$), MV OR = 3.09 ($p = 9.6 \times 10^{-6}$); G1 (24/71): CMH OR = 2.11 ($p = 0.025$), MV OR = 2.58 ($p = 4.0 \times 10^{-3}$) |
| Stage 2 | *RRE* ↔ *rev2* (7880 ↔ 8500) | 72.6% (73.6% G1) | 61.1% | 72.2% / 72.7% (81.7%) | All: CMH OR = 1.91 ($p = 2.1 \times 10^{-3}$), MV $p = 0.027$ (0.119 cond.); G1 (58/71): $p_{\text{lin}} = 0.046$, CMH OR = 2.31 ($p = 0.021$), MV OR = 2.67 ($p = 4.0 \times 10^{-3}$) |
| Stage 2 | *gag/pol* Hinge (1980–2620) | 7.4% (5.2%) | 7.0% | 10.7% / 15.1% (9.3%) | All: CMH OR = 1.73 ($p = 0.0080$); DAG *de novo* (51/527 = 9.7%, $p = 0.042$); attenuates in G1 (9.3%, $p = 0.16$) |
| Stage 2 | *Pol*-RT Tolerant (2650–3450) | 9.3% (7.2/9.5) | 6.7% | 11.7% / 10.8% (9.9%) | Crossover-tolerant: DAG *de novo* 63/527 = 12.0% ($p = 1.2 \times 10^{-4}$ vs. URF); G1 CMH $p = 0.15$ |
| Stage 2 | *gp41*-CT Return Swap (ret.) | — | 35.6% | 19.5% / 16.7% (23.5%) | Confounded: crude $p = 0.0042$ (110/309 vs. 17/87) vanishes under CMH (OR = 0.79, $p = 0.60$) & G1 (8/34, CMH OR = 0.86, $p = 0.97$) |
| Ctrl | *tat1* ↔ *tat2* (5950 ↔ 8400) | 50.3% lin | 52.7% | 56.3% / 54.5% (63.4%) | All: CMH $p = 0.42$, MV $p = 0.16$; G1: CMH $p = 0.16$, MV $p = 0.093$ (ns) |
| Ctrl | *Gag*-MA ↔ *gp41*-CT (950 ↔ 8650) | 43.2% lin | 46.6% | 39.7% / 50.0% | CMH OR = 0.84 ($p = 0.374$) |
| Ctrl | *Gag*-CA/NC ↔ PR (1550 ↔ 2400) | 67.7% lin | 71.1% | 68.2% / 68.2% | CMH OR = 1.11 ($p = 0.655$) |

†For Stage 1/zone rows, Null lists uniform coding share followed in parentheses by the 2%-noise 1-BP / realistic 6%–10%-div 4-BP EGGMAN detectability nulls (1,204 / 4,126 recalled full-screen BPs; diagnostic-SNP nulls: *gp120* 13.3%, *Gag*-CA 4.8%, 2 rigid cores 18.1%, 4 hinges 41.8%, *gag/pol* 5.6%, *Pol*-RT 7.1%). Co-ancestry rows show the 10,000-rep linear permutation null. ‡Exact HXB2 coordinates of the four flexible inter-domain hinges: *gag.MA/CA* linker (1100–1260), *gag/pol* frameshift & PR/RT (1980–2620), accessory hub *pol.IN..vpu* (4950–6350), and *gp41* TM/CT & *nef* boundary (8150–8950). Canonical full-gene checks: *gag.CA* (1186–1878) Est 0.46× ($p = 5.38 \times 10^{-6}$); *env.gp120* (6225–7757) Est 0.63× ($p = 1.50 \times 10^{-6}$). MV logistic models adjust for total breakpoint count and parental trunk pair. Host deduplication invariance (0.4%/0.8%/1.5% across 31,825 genomes): URF Intact 3′ Block = 22.1%/22.0%/22.2% ($p \leq 0.0051$ vs. Est; $p \leq 0.029$ vs. G1).

Table S10: Empirical multi-pathogen benchmark performance using external `PathogenSpec` panels (`--pack`), BV-BRC surveillance streams, and `ChronAeon` `--temporal-guard` ablations.

| Pathogen Pack | Empirical Cohort | $N$ | Time (s) | Specificity | Recall / Calls | Key Operating Finding |
|---|---|---|---|---|---|---|
| IAV (13,136 nt) | Pure `A1`, `A3` founders | 150 | 0.127 | 99.3% exact (0 false) | — | `A1` 75/75, `A3` 74/75 (1 `OutOfPanel`); zero false reassortants |
| IAV (13,136 nt) | 8 H5N1 reassort. genotypes | 950 | 0.127 | 0 cross-genotype err. | 97.58% (927/950) cert | All 8 genotypes certified incl. `C2.1` (95/100); 22 OOP + 1 URF |
| IAV vs. `GenoFLU` | SRA 1,200 labeled (1,100) | 1,200 | 0.127 | 0 cross-genotype err. | 97.82% vs. 100.0% | 98/100 unassg. structured (92% mosaic); 16/16 seg ML trees (69–100% UFB) |
| IAV OOD | Masked `B3.13` + `D1.1` | 650 | 0.089 | 100.0% novel | 98.92% donor set | `GenoFLU`: 100% `Not_assigned` (1.712 s; 19.2× wall / 77× CPU) |
| IAV BV-BRC | Global H5Nx (3,966 ≥ 11.5 kb) | 3,980 | 7.311 | 388 pure / 402 OOP | 2,756 cert / 420 `URF` | 20 constellations (12 `GenoFLU`, 8 EURL/other) |
| SC2 (29,903 nt) | Curated default ($\gamma$ = 7.0, $m$ = 5) | 1,000 | 0.592 | 100.0% T1 pure | 78.5% T1 / 96.4% state | 100.0% exact pure; 57.0% T1 recomb BP / 92.8% T1+T2; 80.7% T2 exact |
| SC2 (29,903 nt) | Curated CoV ($\gamma$ = 3.5, $m$ = 3) | 1,000 | 1.059 | 98.2% T1 pure | 89.6% T1 / 97.3% state | 99.4% exact pure; 81.0% T1 BP; 80.4% T2 (146/157 `XBB` exact T1) vs. `Nextclade` 98.3% (3.424 s) |
| SC2 Span-16 | World Union (basis disc.) | 10,062 | 2.11 | 100% `BA.4`/`BA.3.2` | 100% `XBB` / 93% `XDV` | Basis-discovery cohort: eliminates 70.5% `BA.4` & 91.3% `BA.3.2` pseudo-mosaics |
| SC2 Span-16 | BV-BRC Stream (indep. val.) | 23,376 | 13.44 | 99.9% def / 99.3% CoV | 100% `XBB`/`XDV` | Indep. validation (13,275 pure): 8× pure drop (5.96% → 0.72%) |
| SC2 BV-BRC | Live 100k (77.4k 2025–26) | 100,000 | 113.1–118.3 | 0 waves > 50% mosaic | 99.1–100% `XBB`; 10 `X*` | 13 clusters (1,434): 97.7% in 10 `X*` roots; 3/79 Wuhan-`B` QA |
| Sarbecovirus | Valid whole genomes | 79 | 0.066 | 5/5 clonal basals | 83.8% (62/74) mosaic | Median 5.0 BPs; 4 all-`N` excluded |
| `ChronAeon` SC2 | 32 clades + 134k guard | 134,446 | 0.04–129.0 | 0/0 col & sub viol. | 0.415 / 0.333 yr | 25 direct + 7 `LSD2_Fallback`; 26/29 DAG ordered |
| `ChronAeon` IAV | 15 jobs + 5,830 guard | 5,830 | 0.09–0.41 | 0/0 col & sub viol. | 0.162 / 0.175 yr | 186 → 0 BV-BRC donor fix; `NAm_B313` 2023.87 |
